# Uncyclized xanthommatin is a key ommochrome intermediate in invertebrate coloration

**DOI:** 10.1101/666529

**Authors:** Florent Figon, Thibaut Munsch, Cécile Croix, Marie-Claude Viaud-Massuard, Arnaud Lanoue, Jérôme Casas

## Abstract

Ommochromes are widespread pigments that mediate multiple functions in invertebrates. The two main families of ommochromes are ommatins and ommins, which both originate from the kynurenine pathway but differ in their backbone, thereby in their coloration and function. Despite its broad significance, how the structural diversity of ommochromes arises *in vivo* has remained an open question since their first description. In this study, we combined organic synthesis, analytical chemistry and organelle purification to address this issue. From a set of synthesized ommatins, we derived a fragmentation pattern that helped elucidating the structure of new ommochromes. We identified uncyclized xanthommatin as the elusive biological intermediate that links the kynurenine pathway to the ommatin pathway within ommochromasomes, the ommochrome-producing organelles. Due to its unique structure, we propose that uncyclized xanthommatin functions as a key branching metabolite in the biosynthesis and structural diversification of ommatins and ommins, from insects to cephalopods.

## 1. Introduction

Ommochromes are widespread phenoxazinone pigments of invertebrates. They act as light filters in compound eyes and determine the integumental coloration of a large range of invertebrates (Figon and Casas, 2019). Ommochromes are also of particular interest in applied sciences as their scaffold has been used to design antitumor agents (Bolognese et al., 2002) and, very recently, to manufacture biomimetic color changing electrochromic devices (Kumar et al., 2018). The two major families of natural ommochromes are the yellow-to-red ommatins and the purple ommins that contain a supplemental phenothiazine ring. Ommatins are currently the best described family of ommochromes and occur throughout invertebrates. Ommins are much less characterized, although they are virtually ubiquitous in insects and cephalopods (Needham, 1974; Riddiford and Ajami, 1971). After nearly 80 years and despite their broad significance, the structural and chemical relationships between these two abundant families of ommochromes remain surprisingly mysterious (Figon and Casas, 2019).

The early steps of the biosynthesis of ommochromes in invertebrates cover the oxidation of tryptophan into kynurenines (Fig 1B) (Figon and Casas, 2019), from insects (Linzen, 1974) and spiders (Croucher et al., 2013) to planarians (Stubenhaus et al., 2016) and cephalopods (Williams et al., 2019b). These oxidative steps are homologous to the kynurenine pathway of vertebrates in which the last enzyme, the mitochondria-bound kynurenine 3-monooxygenase, catalyzes the formation of 3-hydoxykynurenine. This *ortho*-aminophenolic amino acid is the currently accepted last common precursor of all known ommochromes, from ommatins to ommins (Fig 1B) (Figon and Casas, 2019). The catabolism of tryptophan then diverges from vertebrates because invertebrates lack the glutarate pathway but possess the ommochrome pathway (Linzen, 1974).

**Fig 1.**
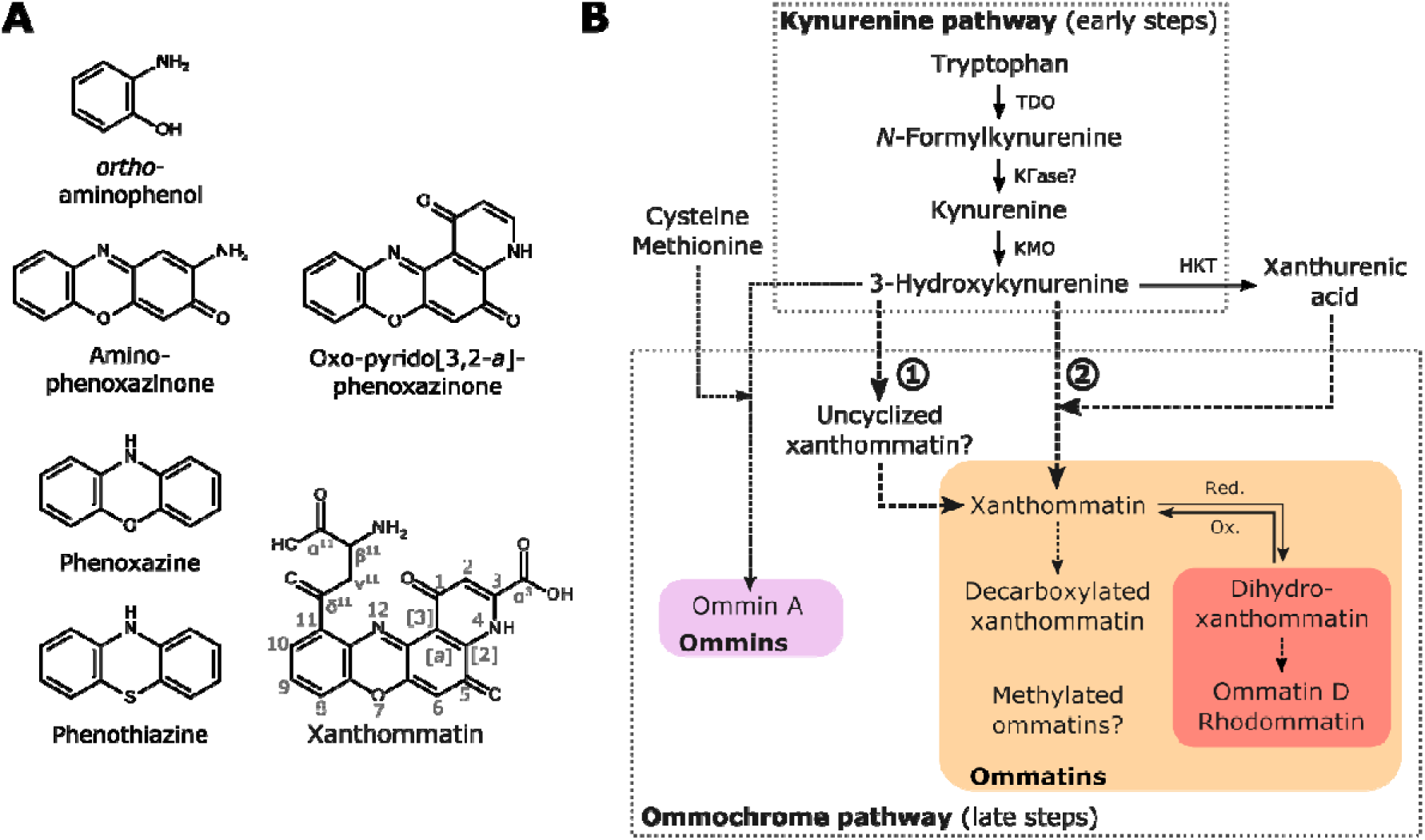
Current knowledge of the tryptophan ommochrome pathway of invertebrates. A) Main chemical structures and chromophores of the tryptophan ommochrome pathway. Numbering of ommatins used in this study is indicated on the structure of xanthommatin. B) Kynurenine and ommochrome pathways form the early and late steps of the tryptophan ommochrome pathway, respectively. Ommatins are possibly biosynthesized via two routes: (1) the dimerization of 3-hydroxykynurenine into the intermediary uncyclized xanthommatin, or (2) the direct condensation of 3-hydroxykynurenine with its cyclized form, xanthurenic acid. Ommatin and ommin pathways share 3-hydroxykynurenine as a precursor, but at which step they diverge is not known. Dashed arrows, steps for which we lack clear biological evidence. HKT, 3-hydroxykynurenine transaminase. KFase, kynurenine formamidase. KMO, kynurenine 3-monooxygenase. Ox., oxidation. Red., reduction. TDO, tryptophan 2,3-dioxygenase.

Ommochromes are produced within specialized intracellular organelles, called ommochromasomes, most likely after the incorporation of 3-hydroxykynurenine (Figon and Casas, 2019; Mackenzie et al., 2000). Two hypotheses have been proposed to explain how ommatins originate from there. (1) It has been suggested very early, but on weak evidence, that the oxidative dimerization of 3-hydroxykynurenine into uncyclized xanthommatin and its subsequent intramolecular cyclization account for the biosynthesis of ommochromes (Fig 1B) (Butenandt and Schäfer, 1962). (2) The condensation of *ortho*-aminophenols with xanthurenic acid has been proposed to form directly the oxo-pyrido[3,2-*a*]phenoxazinone chromophore of ommatins (Fig 1B) (Linzen, 1974; Panettieri et al., 2018). Hypothesis 1 is currently more accepted because ommatins can be synthesized *in vitro* by the oxidative condensation of 3-hydroxykynurenine, which is predicted to form an unstable intermediate, the 3-hydroxykynurenine dimer called uncyclized xanthommatin (Butenandt and Schäfer, 1962; Figon and Casas, 2019; Iwahashi and Ishii, 1997; Williams et al., 2019a; Zhuravlev et al., 2018). Furthermore, uncyclized xanthommatin was speculated in biological extracts (Bolognese and Scherillo, 1974), putatively identified in the *in vitro* oxidation of 3-hydroxykynurenine (Iwahashi and Ishii, 1997) and its enzymatic formation predicted *in silico* by quantum calculations (Zhuravlev et al., 2018). However, it has never been formerly extracted and characterized in biological samples. Alternatively, hypothesis 2 does not involve the formation of any intermediate between the kynurenine pathway and the ommatin pathway. Hence, to discriminate between the two hypotheses, one needs to determine whether uncyclized xanthommatin is produced *in vivo* (Fig 1B). Finding this still-elusive intermediate in biological extracts would therefore be a major step in characterizing the actual biosynthetic pathway of ommochromes.

Deciphering the ommochrome pathway requires the characterization of metabolites in biological extracts. However, biological ommochromes are remarkably refractory to NMR spectroscopy, partly because of their poor solubility in most conventional solvents (Bolognese et al., 1988b; Crescenzi et al., 2004; Parrilli and Bolognese, 1992). It was only very recently that the first ^1^H-NMR spectrum of xanthommatin, whose structure has been known for 60 years, was published after extensive purification and optimization steps (Kumar et al., 2018). However, from a theoretical point of view, mere ^1^H-NMR data cannot provide enough information on the exact structure of ommochromes. Indeed, they are rather poor in carbon-bonded hydrogens, which prevents access to all positions in the structure. Furthermore, they are redox/pH sensitives and prone to tautomerization, which complicate ^1^H spectra greatly. Since our main target compound, uncyclized xanthommatin, is unstable in solution (Bolognese et al., 1988a; Bolognese and Scherillo, 1974), it is highly improbable that gold-standard techniques for structural elucidation, such as ^13^C- and 2D-NMR, can be used because they suffer from too low sensitivity. In order to elucidate the biological diversity of ommochromes, one should therefore look for a combination of more sensitive analytical techniques that provide orthogonal information, such as mass spectrometry and UV-Visible spectroscopy. For nearly a decade, mass spectrometry (MS) has been used to elucidate the structure of both known and unknown ommochromes from biological samples (Futahashi et al., 2012; Panettieri et al., 2018; Reiter et al., 2018; Williams et al., 2016). Yet, evidence for common and compound-specific fragmentation patterns of ommochromes are scarce [but see (Panettieri et al., 2018; Reiter et al., 2018)]. Together with the seldom use of synthesized ommochromes, this lack of analytical data accounts for the very little progress made since four decades to unravel the biological diversity of ommochromes (Figon and Casas, 2019).

In this study, we synthesized xanthommatin and its decarboxylated form by the oxidative dimerization of 3-hydroxykynurenine. Knowing that ommatins are methoxylated in acidified methanol (MeOH-HCl), we incubated synthesized xanthommatin in MeOH-HCl to produce a range of ommatin-derivatives. We constructed an analytical dataset of those ommatins by a combination of UV-Visible spectroscopy, and (tandem) mass spectrometry after separation by liquid chromatography. From this dataset, we derived a fragmentation pattern with valuable structural information, especially when combined with UV-Visible spectra, to infer the structure of new ommochromes with strong confidence. Hence, we could elucidate the structure of three methoxylated ommatins and, more importantly, of uncyclized xanthommatin. Our experiments demonstrated that ommatins are easily and rapidly methoxylated leading to artifacts in conditions matching standard extraction procedures from biological samples. By combining our analytical tools with an artifact-free extraction protocol and a subcellular fractionation of ommochromasomes, we reinvestigated the ommochromes of housefly eyes. We could identify xanthommatin, decarboxylated xanthommatin and uncyclized xanthommatin in ommochromasomes. Our results provide strong support to the hypothesis that ommatin biosynthesis occurs in subcellular organelles through the dimerization of 3-hydroxykynurenine and its subsequent intramolecular cyclization (hypothesis 1, Fig 1B). Furthermore, the unique structure of uncyclized xanthommatin makes it a good candidate to link the biosynthetic pathways of ommatins and ommins, which has important implications on how ommochromes have diversified in a wide range of phylogenetically-distant species.

## 2. Material and Method 2.1 Insects

Houseflies (*Musca domestica*) were obtained at the pupal stage from Kreca. After hatching, houseflies were either directly processed for ommochromasome purification or stored at −20 °C for ommochrome extraction.

### 2.2 Reagents

Sodium dihydrogen phosphate, sodium hydrogen phosphate, L-kynurenine (≥ 98 %), 3-hydroxy-D,L-kynurenine, trifluoroacetic acid (TFA), Triton X-100, tris(hydroxymethyl)aminomethane (Tris), potassium ferricyanide, magnesium chloride, potassium chloride, potassium pentoxide and cinnabarinic acid (≥ 98 %) were purchased from Sigma-Aldrich. Methanol, potassium chloride and hydrochloric acid (37 %) were purchased from Carlo Erba reagents. Nycodenz® was purchased from Axis Shield. L-tryptophan (≥ 99 %) and xanthurenic acid (≥ 96 %) were purchased from Acros Organics. Sucrose (99 %) and sulfurous acid (≥ 6 % SO_2_) were purchased from Alfa Aesar. β-Mercaptoethanol was purchased from BDH Chemicals. Acetonitrile and formic acid were purchased from ThermoFischer Scientific.

### 2.3 In vitro synthesis of xanthommatin

#### 2.3.1 Oxidative condensation of 3-hydroxykynurenine under anoxia

A mixture of ommatins was synthesized by oxidizing 3-hydroxy-D,L-kynurenine with potassium ferricyanide as previously described (Butenandt et al., 1954; Hori and Riddiford, 1981), with some modifications. In a round bottom flask under argon, a solution of 44.6 mM of 3-hydroxy-D,L-mol (102 mg) in 10.2 mL of 0.2 M phosphate buffer at pH 7.1 (PB). In a second round bottom flask under argon, 174 mM of potassium ferricyanide (303 mg) were dissolved in 5.3 mL of PB. Both solutions were purged with argon and protected from light. The potassium ferricyanide solution was added slowly to the solution of 3-hydroxy-D,L-kynurenine. The resulted reaction mixture was stirred at room temperature for 1 h 30 in darkness. Then, 10 mL of sulfurous acid diluted four times in PB was added. The final solution was brought to 4 °C for 30 min during which red flocculants formed. The suspension was then transferred into a 50 mL centrifuge tube. The round bottom flask was rinsed with 8 mL of sulfurous acid previously diluted four times in PB to ensure the complete reduction and flocculation of synthesized ommatins, as well as to remove ferrocyanide. The final suspension was centrifuged for 10 min at 10 000 × g and at 4 °C. The solid was desiccated overnight under vacuum over potassium hydroxide and phosphorus pentoxide. 104 mg of a reddish brown powder was obtained and kept at 4 °C in darkness until further use.

#### 2.3.2 Solubilization and analyses of synthesized ommatins

A solution of synthesized ommatins at 1 mg/mL of was made in methanol acidified with 0.5 % HCl and pre-cooled at −20 °C (MeOH-HCl). The solution was mixed for 30 s and filtered on 0.45 μ filters. All steps were performed, as much as possible, at 4 °C in darkness. The overall procedure took less than two minutes. Immediately after filtration, the solution was subjected to absorption and mass spectrometry analysis (see below). The filtered solution was then stored at 20 °C in darkness and subjected to the same analysis 24 hours later.

### 2.4 Nuclear Magnetic Resonance (NMR) Spectroscopy

One milligram of synthesized product was solubilized in 600 µL of d6-DMSO acidified with 25 µL of TFA, as previously described (Kumar et al., 2018; Williams et al., 2019a). NMR spectra were recorded on Bruker AVANCE AV 300 instruments and the NMR experiment was reported in units, parts per million (ppm), using residual solvent peaks d6-DMSO (δ = 2.50 ppm) for ^1^H NMR as internal reference. Multiplicities are recorded as: s = singlet, d = doublet, t = triplet, dd = doublet of doublets, m = multiplet, bs = broad singlet. Coupling constants (J) are reported in hertz (Hz). ^1^H NMR (300 MHz, d6-DMSO + 0.04 % TFA) δ 8.37 (bs, 3H, H_1_), 8.21 (bs, 1H, NH_8_), 8.04 (t, *J* = 4.5 Hz, 1H, H_5_), 7.83 - 7.82 (m, 2H, H_4_, H_6_), 7.69 (s, 1H, H_7_), 6.67 (s, 1H, H_9_), 4.46 - 4.42 (m, 1H, H_2_), 3.89 (bs, 2H, H_3_).

### 2.5 Ultra-Pressure Liquid Chromatography coupled to Diode-Array Detector and Electrospray Ionization Source-based Mass Spectrometer (UPLC-DAD-ESI-MS) 2.5.1 System

A reversed-phase ACQUITY UPLC® system coupled to a diode-array detector (DAD) and to a Xevo TQD triple quadrupole mass spectrometer (MS) equipped with an electrospray ionization source (ESI) was used (Waters, Milford, MA). Tandem mass spectrometry (MS/MS) was performed by collision-induced dissociation with argon. Data were collected and processed using MassLynx software, version 4.1 (Waters, Milford, MA).

#### 2.5.2 Chromatographic conditions

Analytes were separated on a CSH™ C18 column (2.1 x 150 mm, 1.7 μm) equipped with a CSH™ C18 VanGuard™ pre-column (2.1 x 5 mm). The column temperature was set at 45 °C and the flow rate at 0.4 mL/min. The injection volume was 5 μL. The mobile phase consisted in a mixture of MilliQ water (eluent A) and acetonitrile (eluent B), both prepared with 0.1 % formic acid. The linear gradient was set from 2 % to 40 % B for 18 min.

#### 2.5.3 Spectroscopic conditions

The MS continuously alternated between positive and negative modes every 20 ms. Capillary voltage, sample cone voltage (CV) and collision energy (CE) were set at 2 000 V, 30 V and 3 eV, respectively, for MS conditions. CE was set at 30 eV for tandem MS conditions. Cone and desolvation gas flow rates were set at 30 and 1 000 L/h, respectively. Absorption spectra of analytes were continuously recorded between 200 and 500 nm with a one-nm step. Analytes were annotated and identified according to their retention times, absorbance spectra, mass spectra and tandem mass spectra (Table 1).

**Table 1.** Analytical characteristics of ommatin-related compounds found in vitro and in biological extracts.

| Annotation<br>(Formula, calculated<br>MW) | RT<br>(min) | Absorbance<br>peaks | Monocharged<br>ions<br>(*m/z* loss) | Double-charged<br>ions<br>(*m/z* loss) | MS/MS fragments<br>(*m/z* loss) |
| --- | --- | --- | --- | --- | --- |
| *Detected in synthesized ommatins, crude extracts of housefly eyes and ommochromasome extracts* |  |  |  |  |  |
| 3-Hydroxykynurenine<br>(C<sub>10</sub>H<sub>12</sub>N<sub>2</sub>O<sub>4</sub>, 224.21) | 1.6 | 231; 264;<br>376 | 224.9 [M+H]<sup>+</sup>;<br>207.9 (-17); 161.9<br>(-63); 152.0 (-73) | Not detected | 207.7 (-17); 161.9 (-<br>63) |
| Uncyclized xanthommatin<br>(C<sub>20</sub>H<sub>18</sub>N<sub>4</sub>O<sub>8</sub>, 442.38) | 6.7 | 235; 420-450 | 442.9 [M+H]<sup>+</sup>;<br>425.9 (-17); 408.8<br>(-34); 353.0 (-90) | 213.6 [M-17+2H]<sup>2+</sup> | 425.9 (-17); 409.0 (-<br>34); 390.9 (-52); 363.1<br>(-80); 353.0 (-90);<br>344.9 (-98); 335.1 (-<br>108); 317.0 (-126);<br>307.0 (-136) |
| Xanthommatin<br>(C<sub>20</sub>H<sub>13</sub>N<sub>3</sub>O<sub>8</sub>, 423.33) | 11.8 | 234; 442 | 423.9 [M+H]<sup>+</sup>;<br>406.8 (-17); 377.9<br>(-46); 350.9 (-73) | 212.5 [M+2H]<sup>2+</sup>;<br>189.5 (-23) | 406.3 (-17; -18); 360.7<br>(-63); 350.8 (-73);<br>316.8 (-107); 304.8 (-<br>119); 288.9 (-135) |
| Decarboxylated<br>xanthommatin<br>(C<sub>19</sub>H<sub>13</sub>N<sub>3</sub>O<sub>6</sub>, 379.32) | 9.1 | 234; 442 | 379.9 [M+H]<sup>+</sup>;<br>362.9 (-17); 333.9<br>(-46); 306.9 (-73) | 190.5 [M+2H]<sup>2+</sup>;<br>167.5 (-23) | 362.7 (-17; -18); 333.7<br>(-46); 316.8 (-63);<br>306.8 (-73); 290.9 (-<br>89) |
| *Only detected in synthesized ommatins* |  |  |  |  |  |
| Xanthommatin<br>sulphate/phosphate ester<br>(C<sub>20</sub>H<sub>13</sub>N<sub>3</sub>O<sub>11</sub>S, 503.40;<br>C<sub>20</sub>H<sub>14</sub>N<sub>3</sub>O<sub>11</sub>P, 503.31) | 12.4 | 236; 445 | 503.9 [M+H]<sup>+</sup>;<br>453.6 (-50); 430.5<br>(-73?); 379.8 (-<br>124); 350.7 (-153) | 252.5 [M+2H]<sup>2+</sup>;<br>229.5 (-23) | 487.0 (-17); 440.6 (-<br>63); 430.8 (-73); 422.8<br>(-81); 396.5 (-107?);<br>384.9 (-119) |
| Decarboxylated<br>xanthommatin<br>sulphate/phosphate ester<br>(C<sub>19</sub>H<sub>13</sub>N<sub>3</sub>O<sub>9</sub>S, 459.39;<br>C<sub>19</sub>H<sub>14</sub>N<sub>3</sub>O<sub>9</sub>P, 459.30) | 8.5 | 236; 443 | 459.8 [M+H]<sup>+</sup>;<br>442.8 (-17); 413.8<br>(-46); 386.7 (-73) | 230.4 [M+2H]<sup>2+</sup>;<br>207.4 (-23) | 442.8 (-17); 413.7 (-<br>46); 396.8 (-63); 386.8<br>(-73) |
| *Detected in synthesised ommatins incubated in MeOH-HCl in darkness* |  |  |  |  |  |
| α<sup>3</sup>-Methoxy-xanthommatin<br>(C<sub>21</sub>H<sub>15</sub>N<sub>3</sub>O<sub>8</sub>, 437.36) | 12.6 | 217; 303;<br>452 | 437.9 [M+H]<sup>+</sup>;<br>420.9 (-17); 391.9<br>(-46); 364.9 (-73) | 219.4 [M+2H]<sup>2+</sup>;<br>196.5 (-23) | 420.7 (-17); 391.8 (-<br>46); 374.8 (-63); 364.8<br>(-73); 314.8 (-123);<br>304.8 (-133) |
| α<sup>3</sup>, α<sup>11</sup>-Dimethoxy-xanthommatin<br>(C<sub>22</sub>H<sub>17</sub>N<sub>3</sub>O<sub>8</sub>, 451.39) | 13.0 | 217; 303;<br>452 | 451.9 [M+H]<sup>+</sup>;<br>434.9 (-17); 391.9<br>(-60); 364.9 (-87) | 226.5 [M+2H]<sup>2+</sup>;<br>196.4 (-30) | 434.9 (-17); 374.8 (-<br>77); 364.9 (-87); 314.8<br>(-137); 304.8 (-147) |
| Decarboxylated α<sup>11</sup>-<br>methoxy-xanthommatin<br>(C<sub>20</sub>H<sub>15</sub>N<sub>3</sub>O<sub>6</sub>, 393.35) | 10.1 | 234; 442 | 393.9 [M+H]<sup>+</sup>;<br>376.9 (-17); 333.9<br>(-60); 307.0 (-87) | 197.6 [M+2H]<sup>2+</sup>;<br>167.4 (-30) | 376.7 (-17; -18); 333.8<br>(-60); 316.8 (-77);<br>306.9 (-87); 290.9 (-<br>105) |
| Unknown altered<br>xanthommatin (455) | 14.4 | 242; 442 | 455.9 [M+H]<sup>+</sup> | 228.4 [M+2H]<sup>2+</sup>;<br>205.4 (-23) | 340.1 (-116); 324.6 (-<br>131?); 295.0 (-161);<br>205.2 (-251) |
| Unknown altered methoxy-xanthommatin (469) | 14.9 | 243; 390;<br>452 | 469.8 [M+H]<sup>+</sup> | 235.5 [M+2H]<sup>2+</sup>;<br>212.4 (-23) | 353.7 (-116); 338.8 (-<br>131); 294.7 (-175);<br>204.8 (-265) |
| Unknown altered<br>dimethoxy-xanthommatin<br>(483) | 15.0 | 242; 388;<br>450 | 484.0 [M+H]<sup>+</sup> | 242.5 [M+2H]<sup>2+</sup>;<br>212.5 (-30) | 353.9 (-130); 338.8 (-<br>145); 294.8 (-189);<br>204.9 (-279) |
| *Detected in synthesized ommatins incubated with β-mercaptoethanol in darkness and in ommochromasome extracts* |  |  |  |  |  |
| β-mercaptoethanol-added<br>xanthommatin<br>(C<sub>22</sub>H<sub>17</sub>N<sub>3</sub>O<sub>9</sub>S, 499.45) | 11.7 | 238; 415 | 499.9 [M+H]<sup>+</sup>;<br>426.8 (-73) | 250.4 [M+2H]<sup>2+</sup>;<br>227.5 (-23); 218.4<br>(-32) | 482.7 (-17); 464.7 (-<br>35) |
| β-mercaptoethanol-added<br>decarboxylated<br>xanthommatin<br>(C<sub>21</sub>H<sub>17</sub>N<sub>3</sub>O<sub>7</sub>S, 455.44) | 9.1 | 232; 412 | 455.9 [M+H]<sup>+</sup>;<br>420.0 (-36); 382.8<br>(-73); 343.8 (-112) | 228.4 [M+2H]<sup>2+</sup>;<br>205.4 (-23); 196.5<br>(-32) | 438.8 (-17); 421.0 (-<br>35); 392.7 (-63) |

### 2.6 Thermal reactivity of ommatins in acidified methanol in darkness

#### 2.6.1 Conditions of solubilization and incubation

Solutions (n = 5) of synthesized ommatins at 1 mg/mL were prepared in MeOH-HCl. The solutions m filters. Aliquots of 50 μL were prepared for each sample and stored at either 20 °C or −20 °C in darkness. All steps were performed, as much as possible, in darkness and at 4 °C. The overall procedure for each sample took less than two minutes. During the course of the experiment, each aliquot was analyzed only once by UPLC-ESI-MS/MS, representing a single time point for each sample.

#### 2.6.2 Quantification of ommatins

Unaltered (*i.e.* xanthommatin and its decarboxylated form) and their methoxylated forms were detected and quantified by absorption and MS/MS (single reaction monitoring [MRM] mode) spectrometry. MRM conditions were optimized for each ommatin based on the following parent-to-product ion transitions: xanthommatin [M+H]^+^ 424>361 *m/z* (CV 38 V, CE 25 eV), ^3^-methoxy-α xanthommatin [M+H]^+^ 438>375 *m/z* (CV 37 V, CE 23 eV), α^3^, α^11^-dimethoxy-xanthommatin [M+H]^+^ 452>375 *m/z* (CV 38 V, CE 25 eV), decarboxylated xanthommatin [M+H]^+^ 380>317 *m/z* (CV 34 V, CE 28 eV) and decarboxylated α-methoxy-xanthommatin [M+H] 394>317 *m/z* (CV 34 V, CE 28 eV). See Fig S2 for detailed information on MS/MS optimization. Peak areas for both absorption and MRM signals were calculated by integrating chromatographic peaks with a “Mean” smoothing method (window size: ± 3 scans, number of smooths: 2). Absorbance values at 414 nm of unaltered ommatins were summed and reported as a percentage of the total absorbance of unaltered and methoxylated ommatins. The decay at −20 °C of uncyclized xanthommatin was followed by integrating both the absorbance at 430 nm and the 443 *m/z* SIR signals associated to the chromatographic peak of uncyclized xanthommatin at RT 6.7 min.

### 2.7 Extraction and content analysis of housefly eyes

#### 2.7.1 Biological extractions

Five housefly (*M. domestica*) heads were pooled per sample (n = 5), weighted and homogenized in 1 mL MeOH-HCl with a tissue grinder (four metal balls, 300 strokes/min for 1 min). The obtained crude extracts were centrifuged for 5 min at 10 000 × g and 4 °C. The supernatants were filtered on 0.45 μ filters and immediately processed for absorption and MS analyses. All steps were performed, as much as possible, in darkness at 4 °C. The overall extraction procedure took less than 20 min.

#### 2.7.2 Chromatographic profile

The chromatographic profile of housefly eyes that includes L-tryptophan, xanthurenic acid, 3-D,L-hydroxykynurenine, uncyclized xanthommatin, xanthommatin and decarboxylated xanthommatin is reported based on their optimized MRM signals. L-Tryptophan and xanthurenic acid were quantified based on their optimized MRM signals: [M+H]^+^ 205>118 *m/z* (CV 26 V and CE 25 eV) and [M+H]^+^ 206>132 *m/z* (CV 36 V and CE 28 eV), respectively. See Fig S2 for detailed information on MS/MS optimization. 3-D,L-Hydroxykynurenine and uncyclized xanthommatin were quantified based on their absorption at 370 and 430 nm, respectively. L-Tryptophan, 3-D,L-hydroxykynurenine and xanthurenic acid levels were converted to molar concentrations using calibrated curves of commercial standards. Molar concentrations of uncyclized xanthommatin are reported as cinnabarinic equivalent, since both metabolites possess the same chromophore and presented similar absorbance spectra (see Fig 5C).

### 2.8 Purification and content analysis of ommochromasomes

#### 2.8.1 Isolation buffers

Ommochromasomes from housefly eyes were purified as previously described (Cölln et al., 1981), with some modifications. Isolation buffer (IB) was prepared with 10 mM Tris, 1 mM MgCl_2_, 25 mM KCl and 14 mM β-mercaptoethanol in distilled water. The pH was brought to 7.0 with 1 M HCl. Isolation buffer with sucrose (IB sucrose) was prepared by adding 0.25 M sucrose to IB. IB and IB sucrose were kept at 4 °C no longer than a day to avoid β-mercaptoethanol degradation (Stevens et al., 1983). All steps of the purification procedure were performed at 4 °C.

#### 2.8.2 Homogenization of housefly eyes

A total of 416 fresh housefly heads were homogenized in 12 mL of IB sucrose with a glass potter Elvehjem homogenizer. The suspension was filtered on gauze and the filtrate recovered. The potter was rinsed with 6 mL of IB sucrose, filtered and the filtrate recovered.

#### 2.8.3 Differential centrifugation

The two filtrates were combined and centrifuged for 2 min at 400 × g. The supernatant was recovered. The pellet was resuspended in 6 mL IB sucrose, centrifuged for 2 min at 400 × g and the supernatant recovered. The supernatants were combined and centrifuged for 5 min at 180 × g. The supernatant was recovered and centrifuged for 12 min at 10 000 × g. The obtained pellet was stored overnight at 4 °C. To limit membrane destabilization and loss of pigments by the action of detergents, the pellet was only resuspended in 12 mL IB sucrose with 30 µL Triton X-100 immediately before ultracentrifugation. The suspension containing Triton X-100 was further centrifuged for 1 min at 100 × g and the supernatant recovered.

#### 2.8.4 Ultracentrifugation

The supernatant was layered onto two discontinuous gradients of (from bottom to top): 2.5 M, 2.25 M, 2 M, 1.75 M, 1.5 M, 1.25 M and 1 M sucrose in IB. The tubes were ultracentrifuged for 45 min at 175000 × g in a Beckman LE-70 ultracentrifuge with a SW32 rotor. The obtained pellets were resuspended in 3 mL of IB sucrose, combined and layered onto a discontinuous gradient of (from bottom to top): 0.99 M, 0.83 M and 0.73 M Nycodenz® (iohexol) in IB. The tube was ultracentrifuged for 2 h 40 at 175 000 × g with a SW41Ti rotor. Purified ommochromasomes were recovered from the 0.83 M layer (corresponding density around 1.4) and centrifuged for 10 min at 23 000 × g. The obtained pellet was rinsed with IB sucrose, resuspended in 1 mL IB sucrose and fractionated into 0.1 mL aliquots. Those aliquots were centrifuged for 20 min at 12 000 × g. The supernatants were discarded and one pellet was directly processed for electron microscopy to check for the absence of membrane contaminants and other organelles, particularly mitochondria and lysosomes (Fig 6A). The remaining obtained pellets of purified ommochromasomes were stored at −20 °C until further use.

#### 2.8.5 Extraction of ommochrome-related metabolites from purified ommochromasomes

Pellets (n = 5) were resuspended in 50 μL MeOH-HCl and directly subjected to UPLC-DAD-ESI-MS/MS analysis. All steps were performed, as much as possible, in darkness and at 4 °C. The overall extraction procedure took less than 2 min per sample.

#### 2.8.6 Metabolic analysis of purified ommochromasomes

The chromatographic profile of purified ommochromasomes that includes L-tryptophan, xanthurenic acid, 3-D,L-hydroxykynurenine, uncyclized xanthommatin, xanthommatin, decarboxylated xanthommatin and β-mercaptoethanol-added ommatins is reported based on their optimized MRM signals. L-Tryptophan, 3-D,L-hydroxykynurenine, xanthurenic acid and uncyclized xanthommatin were quantified as described for crude extracts of housefly eyes.

### 2.9 Statistical analysis

Statistical analyses were performed using the R software, version 3.4.1 (www.r-project.org). Statistical threshold was set to 0.05. Statistical analyses are briefly described in the captions of Fig 4 and Fig 5, and detailed results are reported in File S2

## 3. Results

### 3.1 UPLC-DAD-MS/MS structural elucidation of synthesized xanthommatin and its in vitro derivatives

Since xanthommatin is commercially unavailable, we achieved its *in vitro* synthesis by oxidative condensation of 3-hydroxykynurenine under anoxia as previously reported (Butenandt et al., 1954). ^1^H-NMR spectroscopy on the product validated that the main synthesized compound was xanthommatin (see Material and Methods and Fig S1) (Williams et al., 2019a). The synthesized product was then solubilized in methanol acidified with 0.5 % HCl (MeOH-HCl) and analyzed by Liquid Chromatography (LC) coupled to Diode-Array Detection (DAD) and Mass Spectrometry (MS) (Fig 2). The product was solubilized extemporaneously to avoid any chemical degradation before LC-DAD-MS analyses (Fig 2A, B). Chromatograms showed two main peaks corresponding to xanthommatin (retention time [RT]: 11.8 min, [M+H]^+^ at *m/z* 424) and decarboxylated xanthommatin (RT: 9.1 min, [M+H]^+^ at *m/*z 380) (Table 1). Two co-eluting peaks were present in trace amounts at RT 8.5 and 11.9 min and were associated to [M+H]^+^ at *m/*z 460 and [M+H]^+^ at *m/*z 504, respectively.

**Fig 2.**
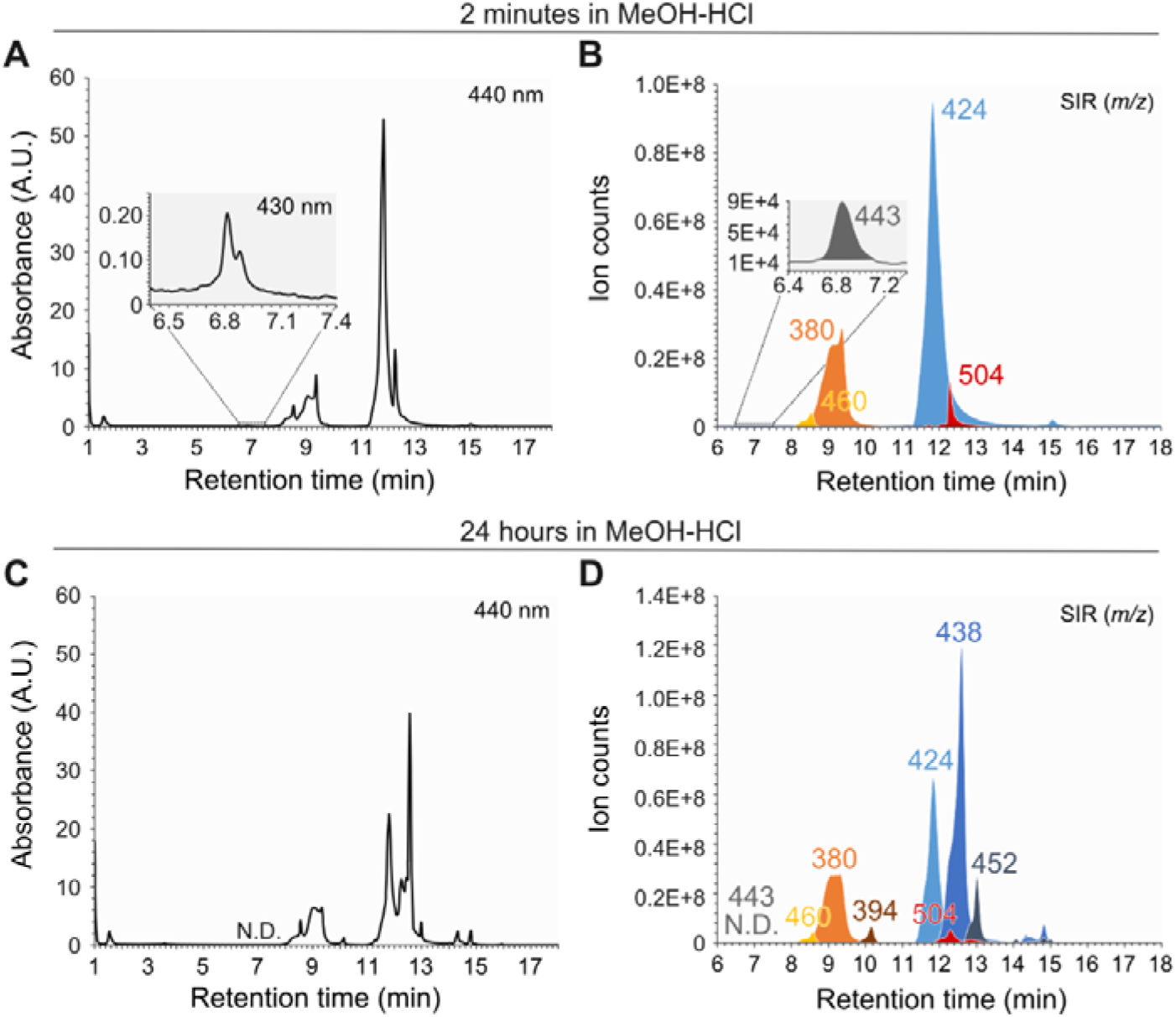
Chromatographic profiles of synthesized xanthommatin before and after storage in acidified methanol. Xanthommatin was synthesized by oxidizing 3-hydroxykynurenine with potassium ferricyanide. **(A-B)** The ommatin solution was subjected to liquid chromatography (LC) two minutes after solubilization in methanol acidified with 0.5 % HCl (MeOH-HCl). The eluted compounds were detected by their absorbance at 440 and 430 nm (A). The main molecular ions (electrospray ionization in positive mode) associated to each peak were monitored by a triple quadrupole mass spectrometer running in single ion reaction (SIR) mode (B). **(C-D)** The same ommatin solution was left for 24 hours at 20 °C in complete darkness. Compounds were separated by LC and detected using the same absorbance (A) and MS modalities (B) as described above.

The detailed analysis of this sample enabled the detection of a small peak at RT 6.7 min ([M+H]^+^ at *m/*z 443) with its corresponding chromatogram at 430 nm (Fig 2A). A detailed analysis of this particular compound is presented further in the text.

With the aim to produce more ommatins and thus manipulate their molecular structure, we incubated synthesized ommatins for 24 h in MeOH-HCl at 20 °C in darkness (Fig 2C, D). Based on previous studies (Bolognese and Liberatore, 1988), we expected ommatins to get methoxylated, mainly on their carboxylic acid functions. The comparative analysis of Fig 2A, B (2 minutes in MeOH-HCl) and Fig 2C, D (24 h later) highlights different sets of peaks that appeared or disappeared over the 24 h of incubation. Three major newly formed compounds were observed at RT 10.1, 12.6 and 13 min corresponding to [M+H]^+^ at *m/*z 394, 438 and 452, respectively. We compared the UV and MS characteristics of these compounds with those of xanthommatin and decarboxylated xanthommatin. Absorbance spectra of the five compounds revealed strong similarities, particularly in the visible region (> 400 nm, Fig 3A) suggesting that these three newly formed molecules shared their chromophores with xanthommatin and decarboxylated xanthommatin. Mass spectra of the five compounds also showed strong similarities (Fig 3B). They all experienced an in-source neutral loss of −17 *m/z* (-NH_3_) and formed a double-charged molecular ion [M+2H]^2+^. 452 and 394 *m/z*-associated compounds typically lost 14 units more than xanthommatin and decarboxylated xanthommatin, respectively, during in-source fragmentation. Additionally, their double-charged fragmentations were 7 units higher than for xanthommatin and decarboxylated xanthommatin (= 14/2; Fig 3B). Overall, gains of 14 *m/z* in the newly formed compounds compared to xanthommatin and decarboxylated xanthommatin suggested methylation reactions occurring in acidic methanol.

**Fig 3.**
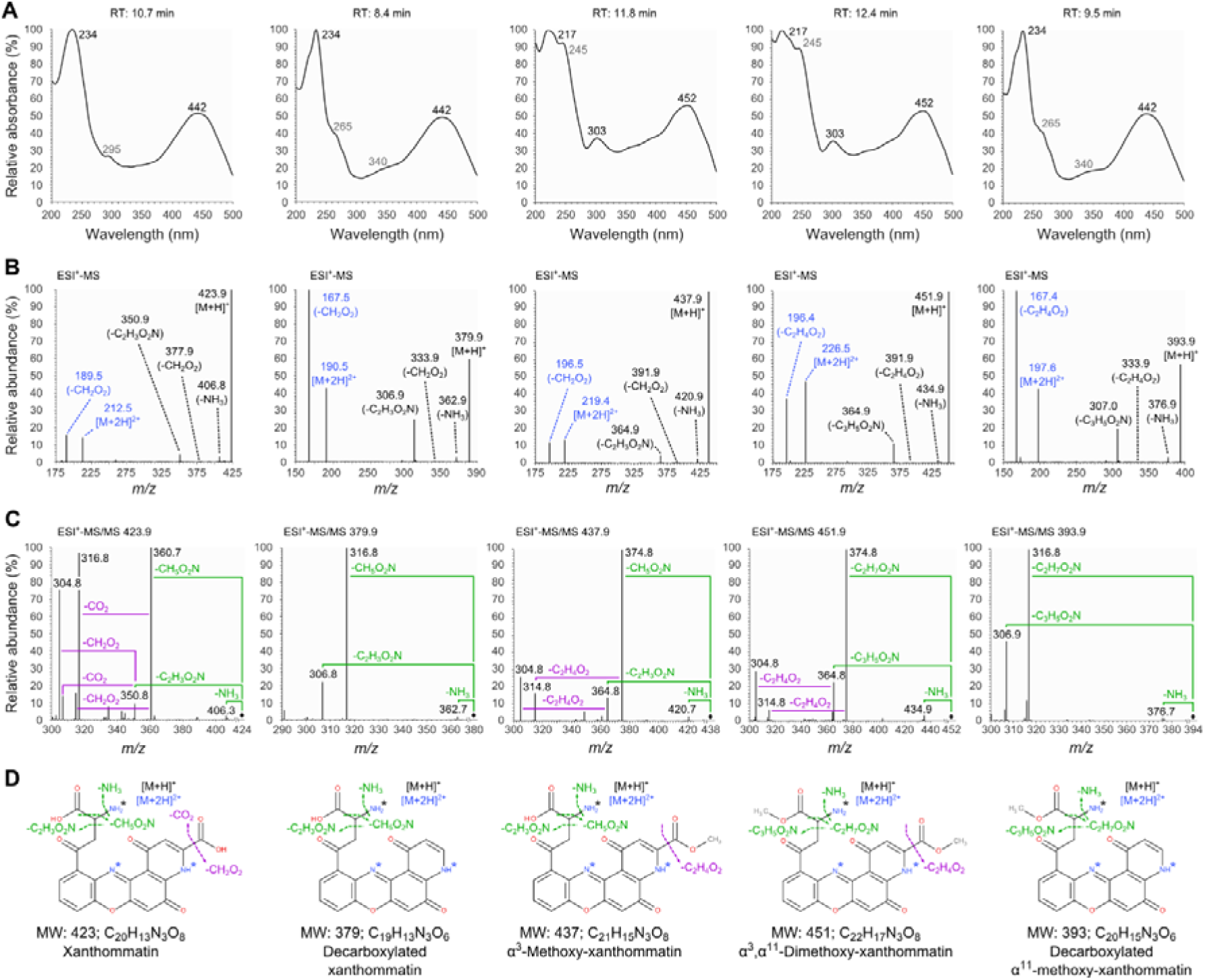
Absorbance- and mass spectrometry-assisted elucidation of the structure of the five major ommatins detected after incubation in acidified methanol. Ommatins incubated for 24 hours in acidified methanol were analysed by liquid chromatography coupled to a photodiode-array detector and a triple quadrupole mass spectrometer. (A) Absorbance spectra. For each metabolite, absorbance values were reported as percentages of the maximum absorbance value recorded in the range of 200 to 500 nm. Major and minor absorbance peaks are indicated in black and grey fonts, respectively. (B) Mass spectra showing molecular ions and in-source fragments. Black fonts, monocharged ions. Blue fonts, double-charged molecular ions. (C) Tandem mass (MS/MS) spectra of molecular ions obtained by collision-induced dissociation with argon. Black diamonds, [M+H]^+^ *m/z*. Green fonts, fragmentations of the amino acid chain. Purple fonts, fragmentations of the pyridine ring are indicated. (D) Elucidated structures of the five ommatins. MS/MS fragmentations are reported in green and purple like in panel C. Black asterisk, main charged basic site. Blue asterisks, potential charged basic sites of the double-charged molecular ions.

Because we did not succeed in purifying each compound to analyze them separately by NMR spectroscopy, we subjected the main [M+H]^+^ to MS/MS fragmentation and compared it with previously reported fragmentation patterns of kynurenine, 3-hydroxykynurenine, xanthommatin and decarboxylated xanthommatin (Guijas et al., 2018; Panettieri et al., 2018; Reiter et al., 2018; Vazquez et al., 2001; Williams et al., 2016) (http://metlin.scripps.edu, METLIN ID: 365). In these molecules, the main ionization site is the amine function of the amino acid branch, which is also the most susceptible to fragmentation. For both xanthommatin and decarboxylated xanthommatin, we observed similar patterns of fragmentation of the amino acid branch with three neutral losses corresponding to - NH_3_ (−17 *m/z*), -CH_5_O_2_N (−63 *m/z*) and -C_2_H_3_O_2_N (−73 *m/z*) (Table 1). Those fragmentations have been reported for these two ommatins (Panettieri et al., 2018; Reiter et al., 2018; Williams et al., 2016), as well as for kynurenines (Vazquez et al., 2001), indicating that they are typical of compounds with a kynurenine-like amino acid chain. Additionally, two neutral losses corresponding to -CO_2_ (−44 *m/z*) and -CH_2_O_2_ (−46 *m/z*) were observed only for xanthommatin (Fig 3C) due to the presence of the carboxyl function on the pyridine ring (Fig 3D). We categorized those predictable MS fragments into different successive fragmentation signatures called F_A_ to F_E_ (Table 2) and we used them to assign the structure of unknown ommatins. The fragmentation of [M+H]^+^ 394 *m/z* showed neutral losses corresponding to -NH_3_ (−17 *m/z* = F_A_), -C_2_H_7_O_2_N (−77 = −63 −14 *m/z* = F_A_ + F_B_ + F_C_ - CH_2_) and - C_3_H_5_O_2_N (−87 = −73 −14 *m/z* = F_A_ + F_B_ + F_C_ + F_D_ - CH_2_) on the amino acid branch. These results strongly indicated that, in the 394 *m/z*-associated compound, the carboxyl function of the amino acid branch was methoxylated (α^11^ position). Consequently, this compound was assigned to decarboxylated α-methoxy-xanthommatin (Fig 3C). Those conclusions are in accordance with the similar absorbance spectra of decarboxylated α-methoxy-xanthommatin and decarboxylated xanthommatin (Fig 3A), as the amino acid branch is unlikely to act on near-UV and visible wavelength absorptions of the chromophore. The fragmentation of [M+H]^+^ 438 *m/z* showed the xanthommatin-like neutral losses, - NH_3_ (−17 *m/z* = F_A_), -CH_5_O_2_N (−63 *m/z* = F_A_ + F_B_ + F_C_) and -C_2_H_3_O_2_N (−73 *m/z* = F_A_ + F_B_ + F_C_ + F_D_), highlighting that this compound shared the same unaltered amino acid branch with xanthommatin.

**Table 2.** Diagnostic neutral losses of ommatins.

| Annotation | Fragmented structure | Neutral loss | $m/z$ loss |
| --- | --- | --- | --- |
| F<sub>A</sub> | Amino acid | -NH<sub>3</sub> | -17 |
| F<sub>B</sub> | Amino acid | -H<sub>2</sub>O | -18 |
| F<sub>C</sub> | Amino acid | -CO | -28 |
| F<sub>D</sub> | Amino acid | -C+2H | -10 |
| F<sub>E</sub> | Pyridine ring | -CO<sub>2</sub>H | -46 |

However, this compound experienced the neutral loss -C_2_H_4_O_2_ (−60 = −46 −14 *m/z* = F_E_ - CH_2_) on the pyridine ring instead of -CH_2_O_2_ (−46 *m/z* = F_E_) (Fig 3C), which strongly indicated a methoxylation on the pyrido-carboxyl group (α^3^ position). This is in accordance with the associated absorbance spectrum being different from that of xanthommatin, which has a carboxylated chromophore (Fig 3A). Hence, we proposed that this compound was α^3^-methoxy-xanthommatin (Fig 3D). The 452 *m/z*-associated compound showed neutral losses corresponding to -NH_3_ (−17 *m/z* = FA), -C_2_H_7_O_2_N (−77 *m/z* = F_A_ + F_B_ + F_C_ - CH_2_) and -C_3_H_5_O_2_N (−87 *m/z* = F_A_ + F_B_ + F_C_ + F_D_ - CH_2_) on the amino acid branch and -C_2_H_4_O_2_ (−60 *m/z* = F_D_ - CH_2_) on the pyridine ring, which strongly indicated methoxylations on both pyridine ring and amino acid branch in positions α^3^ and α^11^, respectively. This is in accordance with the associated absorbance spectrum being similar to that of α^3^-methoxy-xanthommatin, which has a methoxylated chromophore. Thus, we proposed this compound to be α, -dimethoxy-xanthommatin (Fig 3D). Overall, such positions of methoxylation agree well with the reactivity of ommatins in MeOH-HCl (Bolognese and Liberatore, 1988).

Using the same DAD-MS combination approaches, we annotated other ommatin-like compounds produced during *in vitro* synthesis and after incubation in MeOH-HCl (Table 1). The 504 and 460 *m/z*-associated compounds differed from xanthommatin and decarboxylated xanthommatin by 80 Da, respectively. The associated double-charged ions [M+2H]^2+^ and [M-CH_2_O_2_+2H]^2+^ accordingly differed by 40 *m/z* from those of xanthommatin and decarboxylated xanthommatin, respectively. Their absorbance spectra were identical to the two ommatins. Their MS and MS/MS spectra revealed identical losses to xanthommatin and decarboxylated xanthommatin: -NH_3_ (−17 *m/z* = F_A_), -CH_2_O_2_ (−46 *m/z* = F_D_), -CH_5_O_2_N (−63 *m/z* = F_A_ + F_B_ + F_C_) and -C_2_H_3_O_2_N (−73 *m/z* = F_A_ + F_B_ + F_C_ + F_D_; see Table 2). Furthermore, the [M+H]^+^ 504 *m/z* experienced the same neutral losses -C_2_H_5_O_4_N (−107 m/z) and -C_3_H_5_O_4_N (−119 *m/z*) than xanthommatin. Alternatively, the [M+H]^+^ 504 *m/z* experienced a unique in-source neutral loss of −153 *m/z* that could correspond to -C_2_H_3_O_5_NS or -C_2_H_4_O_5_NP (−73 −80 *m/z*). All those results suggested that the 504 and 460 *m/z*-associated compounds were sulphate or phosphate esters of xanthommatin and decarboxylated xanthommatin, respectively. This in accordance with the use of phosphate buffer for the *in vitro* synthesis and of sulfurous acid to precipitate ommatins. To our knowledge, these two esters have never been described. Finally, during the incubation in MeOH-HCl, minor ommatin-like compounds (classified as ommatins based on their absorbance and the presence of double-charged ions) were formed and were associated to the 456, 470 and 484 *m/z* (Table 1). The differences of 14 units of their respective molecular ion *m/z*, as well as their neutral losses in MS/MS being either similar or differing by 14 units, indicated that they were methylated versions of each other. However, due to their very low amounts, we could not unambiguously propose a structure.

These results showed that, in storage conditions mimicking extraction procedures with MeOH-HCl, xanthommatin and its decarboxylated form are methoxylated, even in darkness. Those reactions are likely to result from solvent additions with acidified MeOH-HCl (the most efficient solvent for ommatin extraction). To further characterize the importance of those artifactitious reactions, we followed the kinetic of the five ommatins described in Fig 3D in MeOH-HCl at 20 °C and in darkness.

### 3.2 The ommatin profile is rapidly and readily modified overtime by artifactitious methoxylations in acidified methanol

Because absorbance spectra of all five considered compounds did not differ significantly and because some of them were co-eluted (Fig 2), their detection and quantification were performed by MS/MS in multiple reaction monitoring (MRM) mode. MRM conditions were independently optimized for each compound based on the fragmentation of their amino acid branch (Fig S2).

The MRM signal of xanthommatin rapidly decreased overtime in a linear fashion, with a near 40 % reduction after only one day of incubation (Fig 4A). On the contrary, the MRM kinetics of α - methoxy-xanthommatin had a logarithmic-shape, sharply increasing during the first day before reaching a plateau during the two following days (Fig 4B). Both decarboxylated α-methoxy-xanthommatin and α,-dimethoxy-xanthommatin appeared after a few hours of incubation. Their MRM signal then linearly increased overtime (Fig 4C, E). In parallel, the MRM signal of decarboxylated xanthommatin stayed nearly constant, with only a small increase by 1.13 % over the five first hours (Fig 4D). Those results further validate that xanthommatin was readily methoxylated in darkness, primarily in position α^3^. A slower methoxylation on the amino acid branch could account for the delay in the appearance of the two other methoxylated forms. The levels of decarboxylated xanthommatin did not vary much overtime although its methoxycarbonyl ester was produced (Fig 4D, E). This result could be explained by the concomitant and competitive slow decarboxylation of xanthommatin, a reaction that has already been described in MeOH-HCl upon light radiations (Bolognese et al., 1988b).

Because we cannot compare MRM signal intensities of different molecules, we took benefit of their similar absorption in the visible region, especially at 414 nm (Fig 3A, Fig S3), to quantify their relative amounts. Although less sensitive and less specific than MRM-based detection, the absorbance at 414 nm strongly indicated that, after only one day of incubation, one third of ommatins was methoxylated (Fig 4F). Most of the methoxylated ommatins accumulated during the first 24 hours (Fig 4B). As expected, rates of methoxylation were significantly decreased by incubating synthesized ommatins in MeOH-HCl at −20 °C (Fig S4A, B). After storage of a month at −20 °C, the methoxylated ommatins represented nearly 1.2 % of ommatins (Fig S4C).

**Fig 4.**
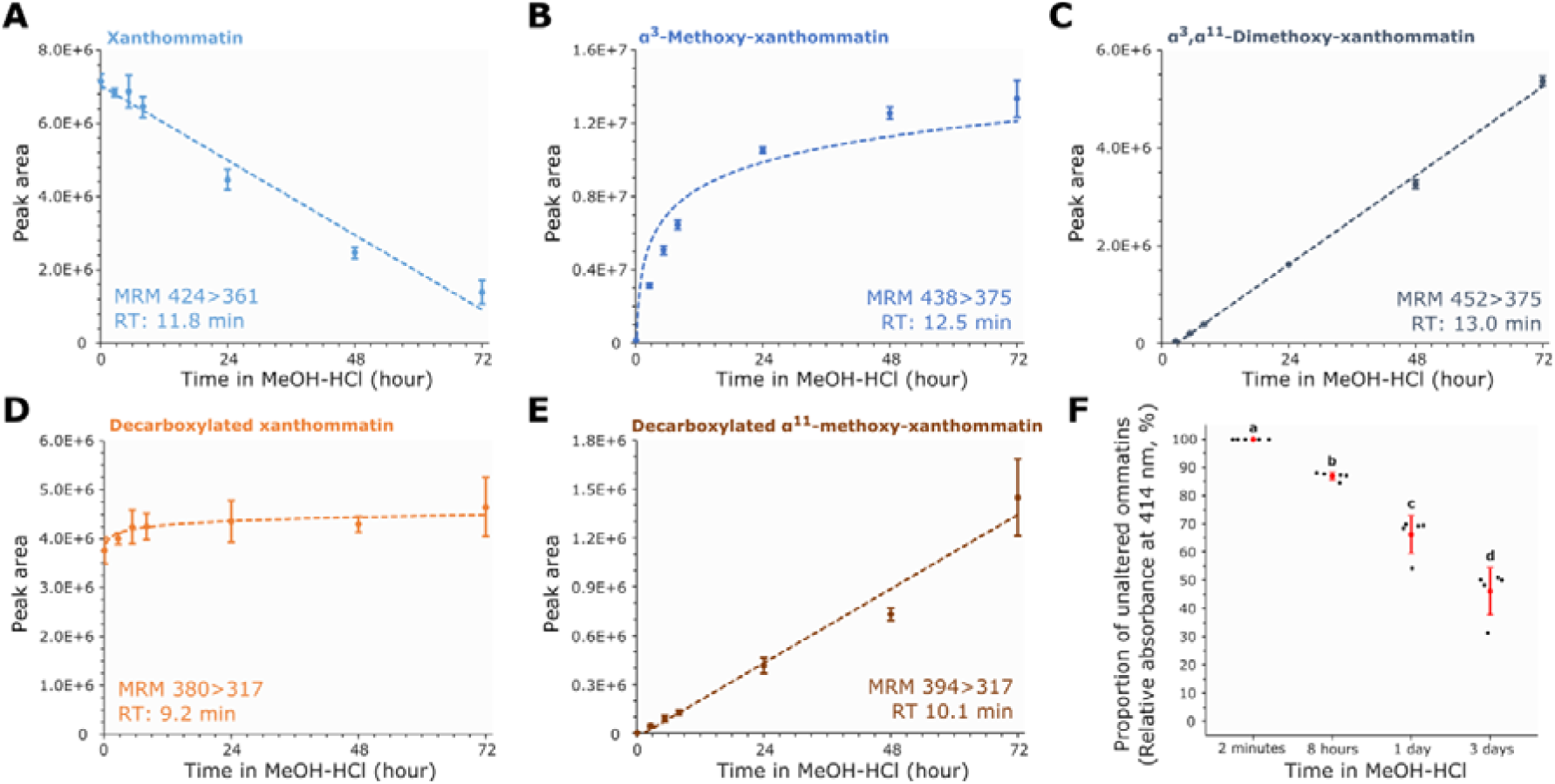
Alterations of synthesized ommatins in acidified methanol at 20 °C in darkness. Synthesized ommatins were solubilized in methanol acidified with 0.5 % HCl (MeOH-HCl) and stored for up to three days at 20 °C and in complete darkness. **(A-E)** Kinetics of alterations were followed by multiple reaction monitoring (MRM) mode of xanthommatin (A), α-methoxy-xanthommatin (B), α, α-dimethoxy-xanthommatin (C), decarboxylated xanthommatin (D) and decarboxylated α-xanthommatin (E). Values are mean ± SD of four to five samples. See Supplemental File S2 for information on regression analyses. **(F)** Relative quantifications of methoxylated ommatins compared to unaltered ones (i.e. xanthommatin and decarboxylated xanthommatin) were performed by measuring the absorbance of ommatins at 414 nm for each time point. Values are mean ± SD of five samples. Different letters indicate statistical differences (Kruskal-Wallis rank sum test: χ = 17.857, df = 3, p-value = 0.00047; pairwise comparisons using Wilcoxon rank sum test and Holm adjustment: p-values < 0.05).

To conclude, our results showed that decarboxylated xanthommatin was mostly stable in MeOH-HCl. By contrast, xanthommatin was rapidly converted into methoxylated derivatives. Since, MeOH-HCl is the most efficient solvent for ommatin extraction, the conditions for extraction and analysis of ommatins from biological samples should avoid wherever possible the formation of artifactitious methoxylated ommatins.

### 3.3 UPLC-DAD-MS/MS structural elucidation of uncyclized xanthommatin, the labile intermediate in the synthesis of ommatins from 3-hydroxykynurenine

The *in vitro* synthesis of xanthommatin by oxidizing 3-hydroxykynurenine additionally yielded a minor compound at RT 6.7 min. It was characterized by a peak of absorbance at 430 nm and was associated to the 443 *m/z* feature (Fig 2A, B). Upon solubilization in MeOH-HCl, the unidentified compound was labile and disappeared after the 24h-incubation at 20 °C in darkness (Fig 2C, D). A similar 443 *m/z* feature was described two decades ago during oxidations of 3-hydroxykynurenine in various conditions (Iwahashi and Ishii, 1997). Based on its MS spectrum, it was assigned putatively to the 3-hydroxykynurenine dimer called uncyclized xanthommatin. However, there was a lack of analytical evidence to support its structural elucidation. No study has ever since reported the presence of uncyclized xanthommatin, either *in vitro* or *in vivo*. Because this compound could be an important biological intermediate in the formation of ommatins (Bolognese et al., 1988b; Iwahashi and Ishii, 1997), we further characterized its structure based on its chemical behavior, absorbance and fragmentation pattern. We note that the apparent lability and the very low amounts of this unidentified product precluded its characterization by NMR spectroscopy.

The absorbance and MS kinetics of the unidentified synthesized compound showed that it was very labile (insets of Fig 5A, B). Indeed, we could not detect it after a week of storage at −20 °C anymore. This behavior resembled that of a photosensitive ommatin-like isolated 40 years ago from several invertebrates, which rapidly turned into xanthommatin after extraction (Bolognese and Scherillo, 1974). The absorbance spectrum of this unidentified compound matched almost exactly the UV-Visible spectrum of cinnabarinic acid measured in the same conditions (Fig 5C), and both are similar to those reported for actinomycin D and 2-amino-phenoxazin-3-one (Nakazawa et al., 1981). This result indicated that this compound contained the amino-phenoxazinone chromophore rather than the pyrido[3,2-*a*]phenoxazinone scaffold of ommatins or the *ortho*-aminophenol core of 3-hydroxykynurenine. Furthermore, its ionization pattern revealed striking similarities with ommatins.

**Fig 5.**
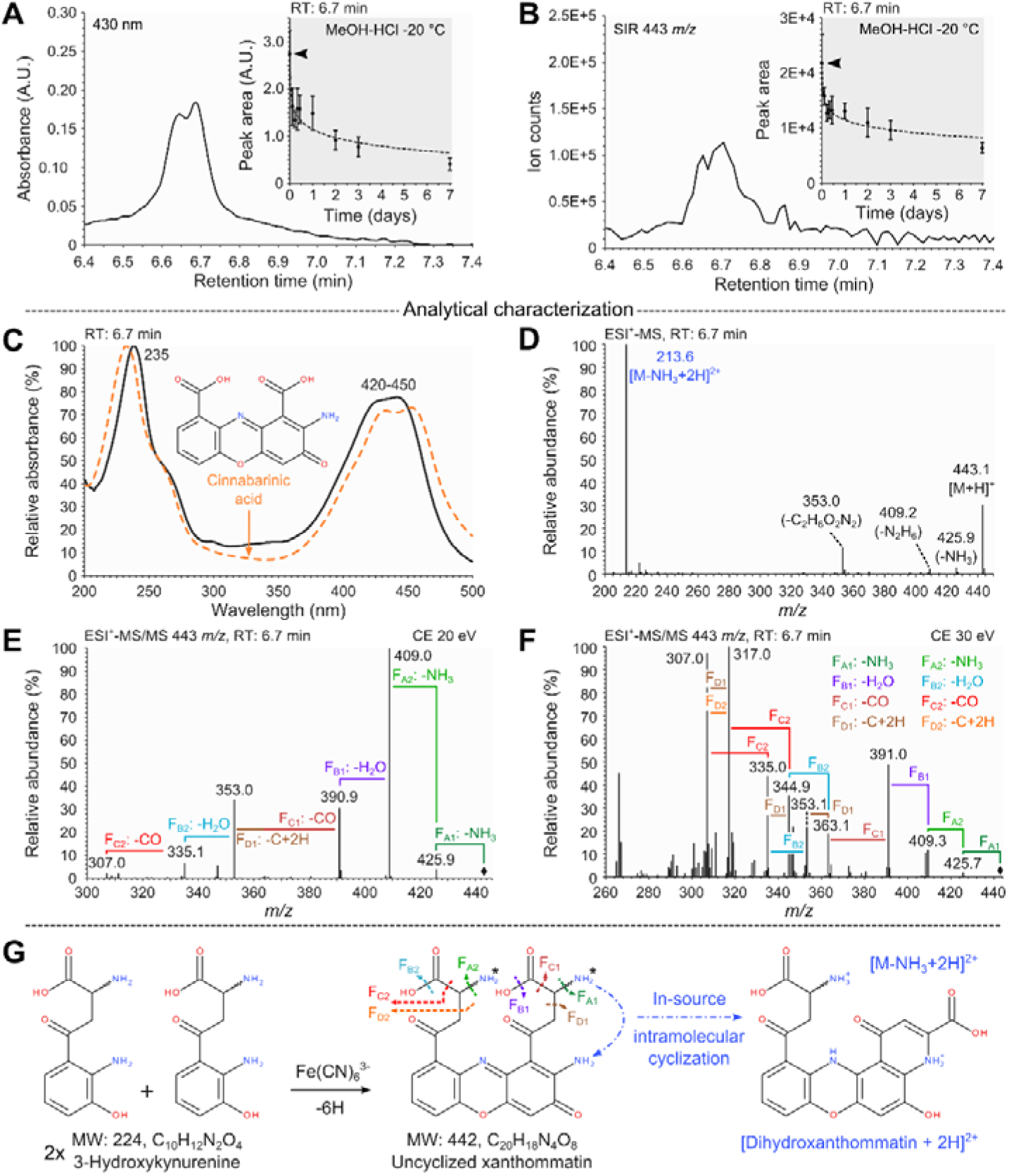
Structural elucidation of uncyclized xanthommatin, the labile intermediate in the synthesis of xanthommatin. **(A-B)** Chromatographic peaks (absorbance at 430 nm [A] and [M+H]^+^ 443 m/z recorded in single ion reaction [SIR] mode [B]) corresponding to the labile ommatin-like compound detected in in vitro synthesis of xanthommatin by the oxidation of 3-hydroxkynurenine with Fe(CN)_6_^3-^. Insets show the decay of chromatographic peaks during storage in methanol acidified with 0.5 % HCl at −20 °C in darkness. Values are mean ± SD of five samples. See Supplemental File S2 for information on regression analysis. **(C)** Absorbance spectra. Solid line, labile ommatin-like compound. Dashed line, the aminophenoxazinone cinnabarinic acid. **(D)** Mass spectrum showing molecular ions and in-source fragments. Black fonts, monocharged ions. Blue font, double-charged ion. **(E-F)** Tandem mass spectra of the molecular ion obtained by collision-induced dissociation with argon at collision energies of 20 eV (E) and 30 eV (F). Fragmentations were classified in four types (F_A_ to F_D_) that occurred twice (F_X1_ and F_X2_). Black diamonds, [M+H]^+^ *m/z*. **(G)** Evidence for the structural elucidation of uncyclized xanthommatin. Colors of the MS/MS fragmentation pattern correspond to those in panels D-F. Black asterisks, potential charged basic sites.

Along with the molecular ion [M+H]^+^ at 443 *m/z* (corresponding to MW 442 and to the formula C_20_H_18_N_4_O_8_), we detected the double-charged ion [M-NH_3_+2H]^2+^ at 213.6 *m/z* (Fig 5D). We then targeted the molecular ion for MS/MS to compare the obtained fragments with those reported above (Table 2). If the compound was uncyclized xanthommatin, we predicted that F_A_, F_B_, F_C_ and F_D_ would appear each twice, because uncyclized xanthommatin possesses two 3-hydroxykynurenine-like amino acid chains. Only F_E_ should be absent, because no aromatic carboxylic acid exist in uncyclized xanthommatin. Indeed, from two MS^2^ spectra obtained at different collision energies, we could assign each F_X_ twice and we did not find any fragmentation event corresponding to F_E_. Hence, by starting from the amino-phenoxazinone backbone suggested by the UV-Visible spectrum, the uncyclized form of xanthommatin could be reconstructed after adding each F_Xx_ successively (Fig 5F).

All these analytical characteristics strongly supported that this labile compound was the phenoxazinone dimer of 3-hydroxykynurenine (Fig 5G), called uncyclized xanthommatin (Figon and Casas, 2019). This structural assignment was further supported by the two following chemical behaviors. First, the oxidation of an *ortho*-aminophenol (here 3-hydroxykynurenine) by potassium ferricyanide is known to induce its dimerization through the loss of six electrons and protons (here 2x MW_3-hydrox_ykynurenine [224 Da] - 6 Da = MWdimer [442 Da] = MW_Uncyclized xanthommatin_). Second, the spontaneous intramolecular cyclization involving the amine functions of the amino-phenoxazinone core and the closest amino acid branch could explain the lability of uncyclized xanthommatin in cold MeOH-HCl (insets of Fig 5A, B) (Williams et al., 2019a), as well as the formation of a major double-charged ion corresponding to that of the reduced form of xanthommatin (dihydroxanthommatin; Fig 5G and Fig S5). In conclusion, we have now the tools to identify uncyclized xanthommatin in other samples, particularly biological materials.

### 3.4 Biological localization of the metabolites from the tryptophan ommochrome pathway

Using our chemical and analytical knowledge of synthesized ommatins, we reinvestigated the content of housefly eyes in ommochromes and their related metabolites. We chose this species because it is known to accumulate xanthommatin and some metabolites of the kynurenine pathway in its eyes (Linzen, 1974). However, nothing is known about decarboxylated xanthommatin and uncyclized xanthommatin because they were recently described [(Figon and Casas, 2019) and this study], even though the presence of uncyclized xanthommatin has been suspected (Bolognese and Scherillo, 1974). Furthermore, a protocol to extract and purify ommochromasomes from housefly eyes is available (Cölln et al., 1981), thus we can address the question of the localization of the metabolites from the tryptophan ommochrome pathway. Finally, we designed an extraction protocol in which all steps were performed in darkness, at low temperature and in less than half an hour. In those conditions, we were confident that artifactitious methoxylated ommatins would represent less than one percent of all ommatins and that we could still detect uncyclized xanthommatin.

Based on MRM signals, we detected in methanolic extractions of housefly eyes (called crude extracts; Fig 6A) the following metabolites of the tryptophan ommochrome pathway: tryptophan, 3-hydroxykynurenine, xanthurenic acid, xanthommatin, decarboxylated xanthommatin and uncyclized xanthommatin (Fig 6B, Table 1). We ascertained the identification of uncyclized xanthommatin by acquiring its absorbance, MS and MS/MS spectra in biological samples. They showed the same features than synthesized uncyclized xanthommatin (Fig S6).

**Fig 6.**
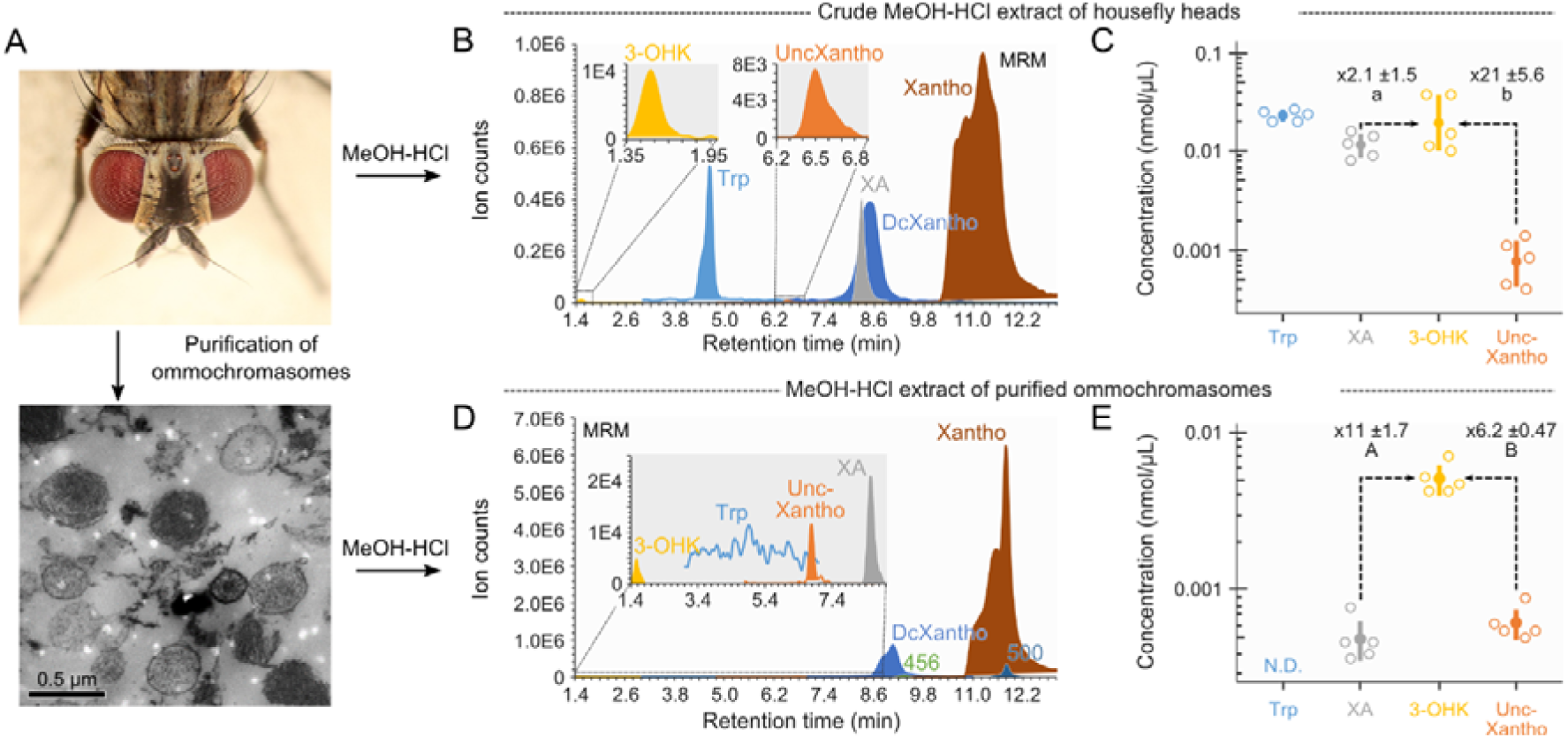
Biological localization of uncyclized xanthommatin and tryptophan ommochrome metabolites from housefly eyes. **(A)** Overview of the purification and extraction protocols of ommochromes from housefly eyes. **(B)** Chromatographic profile in Multiple Reaction Monitoring (MRM) mode of the six main metabolites of the tryptophan ommochrome pathway detected in acidified methanol (MeOH-HCl) extracts of housefly eyes (crude extracts). DcXantho, decarboxylated xanthommatin, 3-OHK, 3-hydroxykynurenine, Trp, tryptophan, UncXantho, uncyclized xanthommatin. Xantho, xanthommatin. XA, xanthurenic acid. **(C)** Five μL of crude extract were injected in the chromatographic system and absolute quantifications of tryptophan, xanthurenic acid, 3-hydroxykynurenine and uncyclized xanthommatin were performed based on available standards (uncyclized xanthommatin levels are expressed as cinnabarinic acid equivalents). Open circles, measures of five independent extracts. Filled circles and error bars, means ± SD of five samples. Ratios of 3-hydroxykynurenine to xanthurenic acid and to uncyclized xanthommatin are shown (mean ± SD, N = 5). **(D-E)** Same as panels B (D) and C (E) but for MeOH-HCl extracts of purified ommochromasomes from housefly eyes. The tryptophan signal was below the signal-to-noise ratio. Statistical differences (p-value < 0.05) between ratios within panels (paired *t*-test) and between panels (unpaired *t*-test) are indicated by different letters and capitals, respectively. N.D., not detected. See Supplemental File S2 for information on statistical analyses. Photograph credits: (A) Sanjay Acharya (CC BY SA).

We then purified ommochromasomes from housefly eyes by a combination of differential centrifugation and ultracentrifugation, and we compared the extracted compounds with those of crude extracts (Fig 6A). The main metabolites of ommochromasomes were xanthommatin and its decarboxylated form (Fig 6D), in accordance to the function of ommochromasomes as ommochrome factories (Figon and Casas, 2019). Based on the absorbance at 414 nm, decarboxylated xanthommatin represented 5.3 ± 0.1 % (mean ± SD, n = 5) of all ommatins detected in extracts of ommochromasomes. In comparison, decarboxylated ommatins represented 21.5 ± 0.2 % (mean ± SD, n = 10) of all ommatins synthesized *in vitro*, a percentage four times higher than in methanolic extracts of purified ommochromasomes (Welch two sample *t*-test, t = 219.99, df = 12.499, p-value < 2.2e^-16^). Regarding the precursors of ommochromes in housefly eyes, we could only detect tryptophan and 3-hydroxykynurenine but not the intermediary kynurenine. Tryptophan remained undetectable in extracts of ommochromasomes (Fig 6D). Xanthurenic acid, the product of 3-hydroxykynurenine transamination, was particularly present in crude extracts of housefly eyes but much less in ommochromasomes relatively to 3-hydroxykynurenine (Fig 6C vs. Fig 6E). We also detected the uncyclized form of xanthommatin in ommochromasomes (Fig 6D). Its level was similar to that of the more stable xanthurenic acid and was enriched relatively to 3-hydroxykynurenine in ommochromasomes compared to crude extracts (Fig 6C vs. Fig 6E).

We detected in ommochromasomes two minor ommatin-like compounds associated to the molecular ions 500 and 456 *m/*z (Table 1), which co-eluted with xanthommatin and its decarboxylated form, respectively (Fig 6D). Both unknown compounds were undetectable in crude extracts of housefly eyes. Their associated *m/z* features differed from those of xanthommatin and decarboxylated xanthommatin by 76 units, respectively (Table 1). Because the isolation buffer used for ommochromasome purifications contained β-mercaptoethanol (MW 78 Da), we tested whether those unknown ommatins could be produced by incubating synthesized ommatins with β-mercaptoethanol in a water-based buffer. We did find that 456 and 500 *m/z*-associated compounds were rapidly formed in those *in vitro* conditions and that their retention times matched those detected in ommochromasome extracts (Fig S7). Therefore, the 456 and 500 *m/z*-associated ommatins detected in ommochromasome extracts were likely artifacts arising from the purification procedure via the addition of β-mercaptoethanol. These results further demonstrate that ommochromes are likely to be altered during extraction and purification procedures, a chemical behavior that should be controlled by using synthesized ommochromes incubated in similar conditions than biological samples.

## 4. Discussion

### 4.1 UPLC-DAD-MS/MS structural elucidation of new ommatins

Biological ommochromes are difficult compounds to analyze by NMR spectroscopy (Bolognese et al., 1988b). Only xanthommatin, the most common and studied ommochrome, has been successfully subjected to ^1^H-NMR (although no ^13^C- and 2D-NMR data exist to date, to the best of our knowledge) (Kumar et al., 2018; Williams et al., 2019a). The NMR-assisted structural elucidation of unknown ommochromes remains therefore extremely challenging and deceptively difficult. To tackle this structural problem, we used a combination of absorption and mass spectroscopies, which offer high sensitivities and orthogonal information, after separation by liquid chromatography. We report here the most comprehensive analytical dataset to date of ommatins, based on their absorbance, mass and tandem mass spectra (Table 1). This dataset allowed us to elucidate with strong confidence the structure of four new ommochromes, including three methoxylated forms and one labile biological intermediate, and to propose structures for eight new other ommatins.

Studies reporting the MS/MS spectra of ommatins demonstrated that they primarily fragment on their amino acid chain (Williams et al., 2016) and then on the pyrido-carboxylic acid, if present (Panettieri et al., 2018). We confirmed those results for other ommatins, which indicates that ommatins fragment in a predictable way despite being highly aromatic compounds. We took a step further by positioning methoxylations based on the differences in fragmentation between methoxylated ommatins and (decarboxylated) xanthommatin. Besides, xanthommatin not only formed a 307 *m/z* fragment but also a major 305 *m/z* fragment (Fig 3C; Table 1). This sole fragment was used in a previous study to annotate a putative new ommochrome called (iso-)elymniommatin, an isomer of xanthommatin (Panettieri et al., 2018). Our results prove that MS/MS spectra cannot distinguish (iso-)elymniommatin and xanthommatin unambiguously. The use of synthesized ommatins and further experiments are thus needed to verify the existence of (iso-)elymniommatin in butterfly wings.

Overall, this analytical dataset will help future studies to identify known biosynthesized and artifactitious ommatins in biological samples, as well as to elucidate the structure of unknown ommatins by analyzing their absorbance and mass spectra in the absence of NMR data. Furthermore, it is now possible to look for uncyclized xanthommatin in a wide variety of species.

### 4.2 Biological extracts are prone to yield artifactitious ommatins

It has long been reported that ommatins are photosensitive compounds that react with acidified methanol (MeOH-HCl) upon light radiation, leading to their reduction, methylation, methoxylation, decarboxylation and deamination (Bolognese et al., 1988c, 1988d; Bolognese and Liberatore, 1988; Figon and Casas, 2019). Nevertheless, incubating tissues in MeOH-HCl for several hours at room temperature has been commonly used to extract ommatins from biological samples efficiently (Bolognese et al., 1988a; Riou and Christidès, 2010; Zhang et al., 2017). Our results demonstrate that, even in the absence of light radiation, ommatins are readily and rapidly methoxylated by thermal additions of methanol, primarily on the carboxylic acid function of the pyridine ring and secondarily on the amino acid chain (Fig 7). The [M+H]^+^ 438 *m/z* of α^3^-methoxy-xanthommatin identified in our study could correspond to the same [M+H]^+^ previously reported in extracts of butterfly wings (Panettieri et al., 2018). Hence, artifactitious methoxylations during extraction should first be ruled out before assigning methoxylated ommatins to a new biosynthetic pathway. We also show that ommatins react with other extraction buffers since we detected β-mercaptoethanol-added ommatins when synthesized ommatins were incubated in a phosphate buffer containing that reducing agent. Overall, these results emphasize the need to control for potential artifactitious reactions when performing any extraction or purification protocol of biological ommochromes.

**Fig 7.**
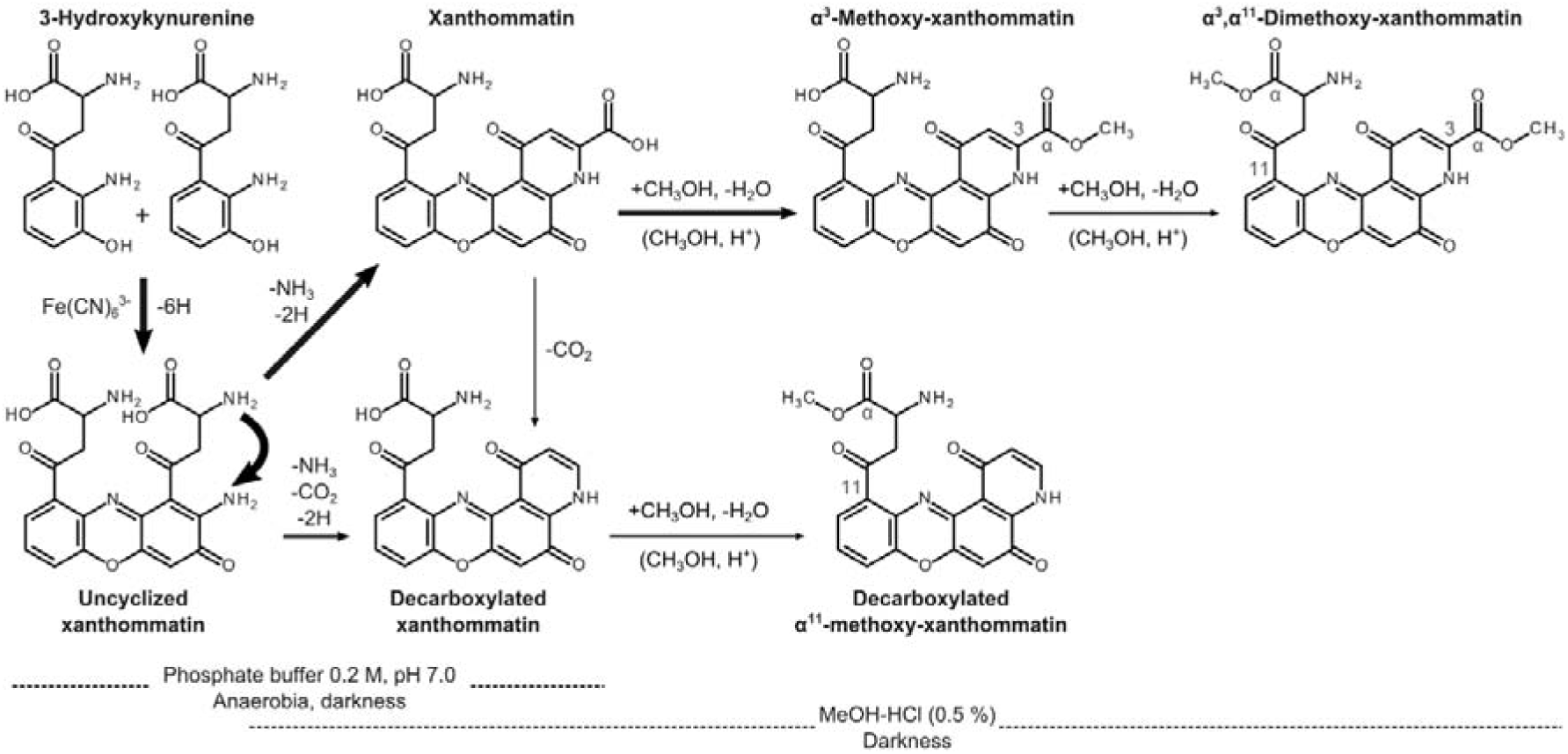
*In vitro* formation and alteration of ommatins. Oxidative condensation of the *ortho*-aminophenol 3-hydroxykynurenine proceeds through the loss of six electrons, leading to the formation of the amino-phenoxazinone uncyclized xanthommatin. Uncyclized xanthommatin then rapidly undergoes intramolecular cyclization and oxidation, forming the two pyrido[3,2-*a*]phenoxazinone xanthommatin and decarboxylated xanthommatin. Decarboxylated xanthommatin could also be produced from the direct decarboxylation of xanthommatin. In acidified conditions, ommatins readily undergo thermal additions of methanol, which leads to their methoxylation. In solution, uncyclized xanthommatin also decays by intramolecular cyclization and xanthommatin slowly decarboxylates. Relative sizes of arrows are indicative of reaction rates.

### 4.3 The metabolites of the tryptophan ommochrome pathway in ommochromasomes

It has long been hypothesized that precursors of ommochromes are translocated within ommochromasomes by the transmembrane ABC transporters White and Scarlet (Ewart et al., 1994; Mackenzie et al., 2000). Here, we clearly demonstrate that 3-hydroxykynurenine, but not tryptophan, occurs in ommochromasome fractions of housefly eyes, confirming that 3-hydroxykynurine is the precursor imported into ommochromasomes by White and Scarlet transporters (Fig 8).

**Fig 8.**
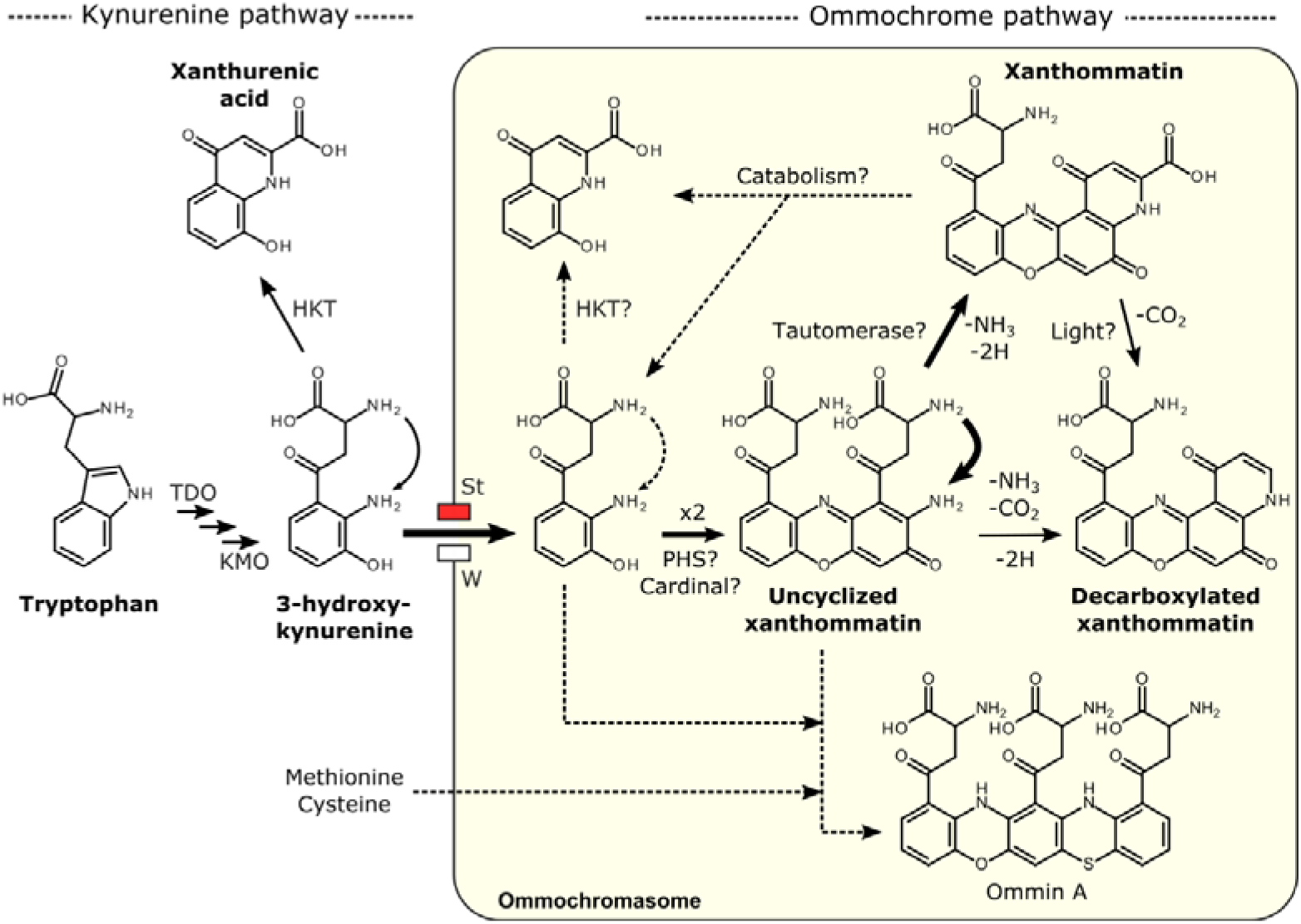
Proposed biosynthetic pathway of ommochromes through the formation of uncyclized xanthommatin in ommochromasomes. See text for more details on each step. Relative sizes of arrows are indicative of reaction rates. TDO, tryptophan 2,3-dioxygenase. KMO, kynurenine 3-monooxygenase. St, ABC transporter scarlet. W, ABC transporter white. PHS, phenoxazinone synthase. HKT, 3-hydroxykynurenine transaminase.

Our results confirm that xanthommatin is the main ommatin in ommochromasomes of housefly eyes. We also showed that decarboxylated xanthommatin was present in significant amounts, which could not be solely due to the slow decarboxylation of xanthommatin in MeOH-HCl. This result indicates that both xanthommatin and its decarboxylated form are produced from 3-hydroxykynurenine within ommochromasomes (Fig 8). We also detected xanthurenic acid, the cyclized form of 3-hydroxykynurenine, in housefly eyes. Compared to 3-hydroxykynurenine, xanthurenic acid was present in minute amounts within ommochromasomes. We hypothesize that xanthurenic acid is produced within ommochromasomes via two non-exclusive pathways (Fig 8). The first route is the *in situ* intramolecular cyclization of 3-hydroxykynurenine, which requires a cytosolic transaminase activity (HKT; Fig 8) (Han et al., 2007). The second route is the degradation of xanthommatin that would produce 3-hydroxykynurenine and xanthurenic acid. The phenoxazinone structure of ommatins is indeed known to undergo ring-cleavage, particularly in slightly basic water-based buffers (Butenandt and Schäfer, 1962). Hence, traces of xanthurenic acid might either come from degradation during the purification protocol or from biological changes in ommochromasome conditions (enzymatic activities or basification) leading to the cleavage of xanthommatin. To the best of our knowledge, no biological pathways for the degradation of the pyrido-phenoxazinone structure of ommatins have been described. The detection of xanthurenic acid in ommochromasomes might therefore be the first step towards understanding the *in situ* catabolism of ommatins (Fig 8).

Experimental and computational chemists have long hypothesized that the pyrido[3,2-*a*]phenoxazinone structure of ommatins should be synthesized *in vivo* by the dimerization of 3-hydroxykynurenine and a subsequent spontaneous intramolecular cyclization (Butenandt, 1957; Zhuravlev et al., 2018). However, the associated dimer of 3-hydroxykynurenine, called uncyclized xanthommatin, proved to be difficult to characterize and to isolate from biological samples because of its lability (Bolognese and Scherillo, 1974). In our study, we synthesized uncyclized xanthommatin by the oxidative condensation of 3-hydroxykynurenine with potassium ferricyanide, an oxidant known to form amino-phenoxazinones from *ortho*-aminophenols (Bolognese and Scherillo, 1974). We used a combination of kinetics and analytical spectroscopy (DAD, MS and MS/MS) to confirm the *in vitro* and biological occurrence of uncyclized xanthommatin. Because we detected xanthommatin, its decarboxylated form, their precursor 3-hydroxykynurenine and the intermediary uncyclized xanthommatin in both *in vitro* and biological samples, we argue that the *in vitro* synthesis and the biosynthesis of ommatins proceed through a similar mechanism (compare Fig 7 and Fig 8). An alternative biosynthetic pathway for ommatins has been proposed to occur through the condensation of 3-hydroxykynurenine with xanthurenic acid (hypothesis 2, Fig 1B) (Linzen, 1974; Panettieri et al., 2018). However, our data do not support this hypothesis because xanthurenic acid, which is a stable compound unlike uncyclized xanthommatin, was present in minute amounts within ommochromasomes compared to 3-hydroxykynurenine, with which it would condensate. At the very least, our results show that xanthurenic acid is tightly linked to the ommochrome pathway and therefore cannot be considered as a marker of a distinct biogenic pathway. Lastly, as far as we know, there has been no experimental evidence for the formation of xanthommatin by condensing 3-hydroxykynurenine with xanthurenic acid. In conclusion, the formation of uncyclized xanthommatin by the oxidative dimerization of 3-hydroxykynurenine is likely to be the main biological route for the biosynthesis of ommatins within ommochromasomes (Fig 8; see Supplemental File S3 for a discussion on the putative enzymatic activity involved in uncyclized xanthommatin formation).

### 4.4 Uncyclized xanthommatin is a potential key branching point in the biogenesis of ommatins and ommins

The relatively recent description of decarboxylated xanthommatin in several species indicates that it is a common biological ommatin (Figon and Casas, 2019). Yet, little is known about how decarboxylation of ommatins proceeds *in vivo*. In this study, we show that decarboxylated xanthommatin is unlikely to arise solely from the artifactitious decarboxylation of xanthommatin in MeOH-HCl, and that the level of decarboxylated xanthommatin is lower in biological extracts (5.3 %) than *in vitro* (21.5 %). Several biological mechanisms could account for the biosynthesis of decarboxylated xanthommatin but we only discuss the one in direct connection with uncyclized xanthommatin (but see Fig 8 and Supplemental File S3 for a discussion of the others). Bolognese and colleagues proposed that decarboxylation happens by a rearrangement of protons, consecutively to the intramolecular cyclization of uncyclized xanthommatin (Bolognese et al., 1988b). Such mechanism has been well described for the biogenesis of eumelanin monomers, in which the non-decarboxylative rearrangement of dopachrome is favored by the dopachrome tautomerase (Solano et al., 1996). Hence, this analogy raises the intriguing possibility that a tautomerase might catalyze the formation of xanthommatin from uncyclized xanthommatin (Fig 8), thereby controlling the relative content of decarboxylated xanthommatin in ommochromasomes, which is known to vary among species, individuals and chromatophores (Futahashi et al., 2012; Williams et al., 2016; Zhang et al., 2017). Why decarboxylated xanthommatin levels depend on the biological context may rely on its biological functions, which we further discuss in the Supplemental File S3.

How ommatins and ommins, the two most abundant families of ommochromes, are biochemically connected to each other is still a mystery (Figon and Casas, 2019). Purple ommins have higher molecular weights than ommatins and derive from both 3-hydroxykynurenine and cysteine/methionine, the latter providing sulfur to the phenothiazine ring of ommins (Linzen, 1974; Needham, 1974). The best-known ommin is called ommin A, whose structure was proposed to be a trimer of 3-hydroxykynurenine in which one of the phenoxazine ring is replaced by phenothiazine (Fig 8) (Needham, 1974). Since the pyrido[3,2-*a*]phenoxazinone cannot be reopen to an amino-phenoxazinone in anyway (Bolognese and Liberatore, 1988), it is unlikely that the biosynthesis of ommins is a side-branch of ommatins. Thus, the biochemical relationship between ommatins and ommins should be found upstream in the biosynthetic pathway of ommochromes. Older genetic and chemical studies demonstrated that ommins and ommatins share the kynurenine pathway and Linzen proposed that the ratio of xanthommatin to ommins could depend on the level of methionine-derived precursors (Linzen, 1974). The distinct structure of uncyclized xanthommatin raises the interesting hypothesis that uncyclized xanthommatin is the elusive intermediate between the *ortho*-aminophenol structure of 3-hydroxykynurenine, the pyrido-phenoxazinone chromophore of xanthommatin and the phenoxazine-phenothiazine structure of ommins. We propose that the biosynthesis of ommins first proceeds with the dimerization of 3-hydroxykynurenine into uncyclized xanthommatin, then with the stabilization of its amino acid chain to avoid a spontaneous intramolecular cyclization, and finally with the condensation with a sulfur-containing compound derived from methionine/cysteine (Fig 8). Although this mechanism is hypothetical at this stage, it can explain two apparently unrelated observations. First, it clarifies the reason why *cardinal* mutants of insects, which lack the heme peroxidase Cardinal that possibly catalyzes the formation of uncyclized xanthommatin, lack both ommatins and ommins, and accumulate 3-hydroxykynurenine (Howells et al., 1977; Osanai-Futahashi et al., 2016). Second, it could explain how a single cephalopod chromatophore can change its color from yellow (ommatins) to purple (ommins) across its lifetime (Reiter et al., 2018). The biochemical mechanism might be analogous to the casing model of melanins (Ito and Wakamatsu, 2008), in which pheomelanins and eumelanins (in this case ommins and ommatins, respectively) are produced sequentially from the same precursors (uncyclized xanthommatin) through changes in sulfur (methionine/cysteine) availability within melanosomes (ommochromasomes).

Overall, uncyclized xanthommatin appears as a key metabolite in the ommochrome pathway by leading to either ommatins, decarboxylated ommatins or ommins (Fig 8). Therefore, the formation of uncyclized xanthommatin might represent a key step in the divergence between the post-kynurenine pathways of vertebrates and invertebrates, as well as in the structural diversification of ommochromes in phylogenetically-distant invertebrates.

## Acknowledgments

We thank Kévin Billet, Cédric Delevoye and Emmanuel Gaquerel for fruitful discussions. We are grateful to Antoine Touzé for his technical assistance and for access to the ultracentrifuge. We thank Rustem Uzbekov for providing the electron micrograph. The ENS de Lyon is thanked for financial support (to F. F.). This study formed part of the doctoral dissertation of F. F. under the supervision of J. C.

## Conflict of interest

The authors declare that they have no conflicts of interest with the contents of this article.

## Abbreviations

CE: collision energy
CV: cone voltage
DAD: diode-array detector
ESI^+^: positive-mode electron spray ionization
LC: liquid chromatography
MeOH-HCl: acidified methanol with 0.5% hydrochloric acid
MRM: multiple reaction monitoring
MS: mass spectrometry
MS/MS: tandem mass spectrometry
MW: molecular weight
*m/z*: mass-to-charge ratio
NMR: nuclear magnetic resonance
RT: retention time
SD: standard deviation
SE: standard error
SIR: single ion recording
UV: ultraviolet

## Supplemental Data

**Fig S1.**
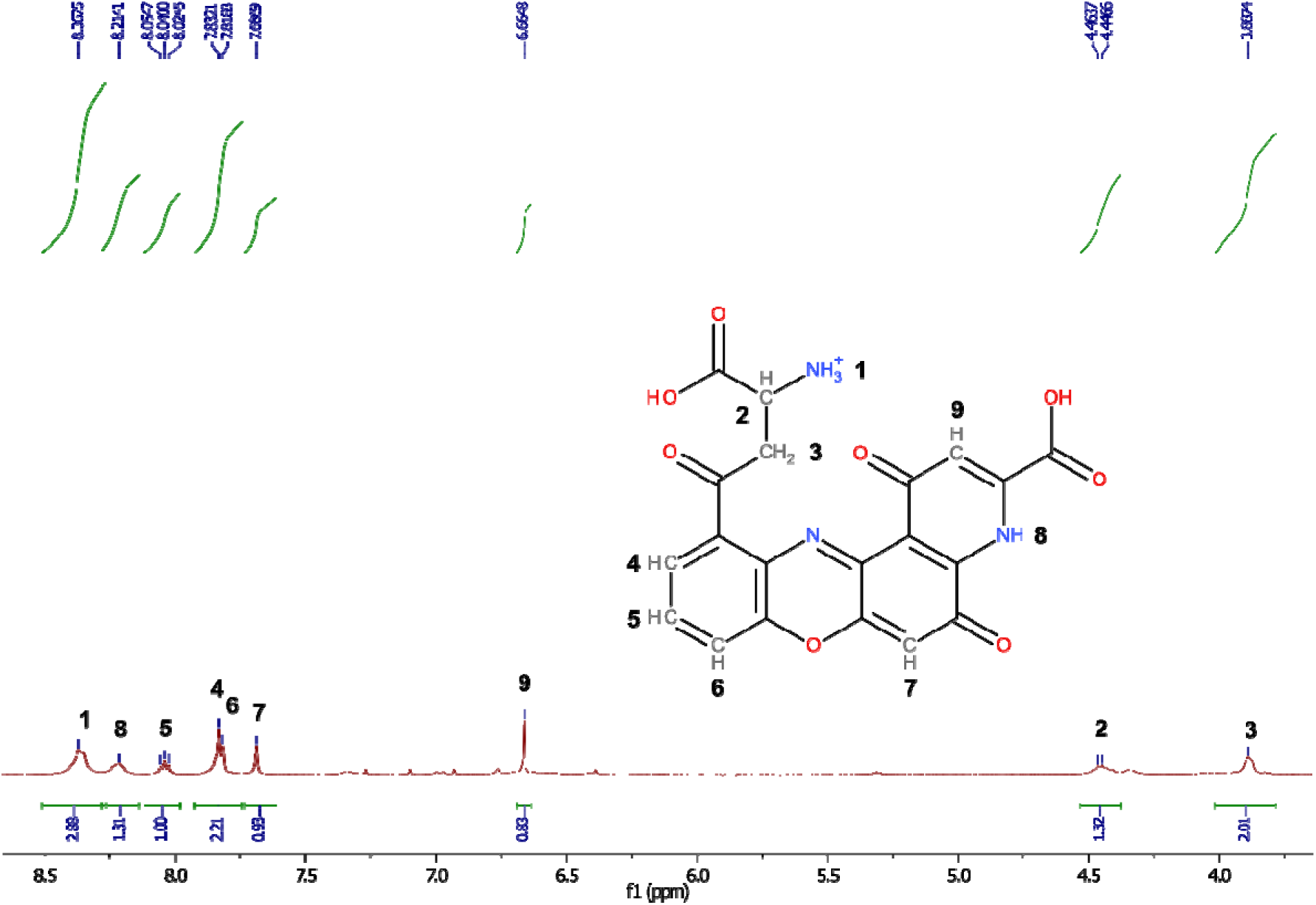
^1^H-NMR spectrum of synthesized xanthommatin in acidic DMSO.

**Fig S2.**
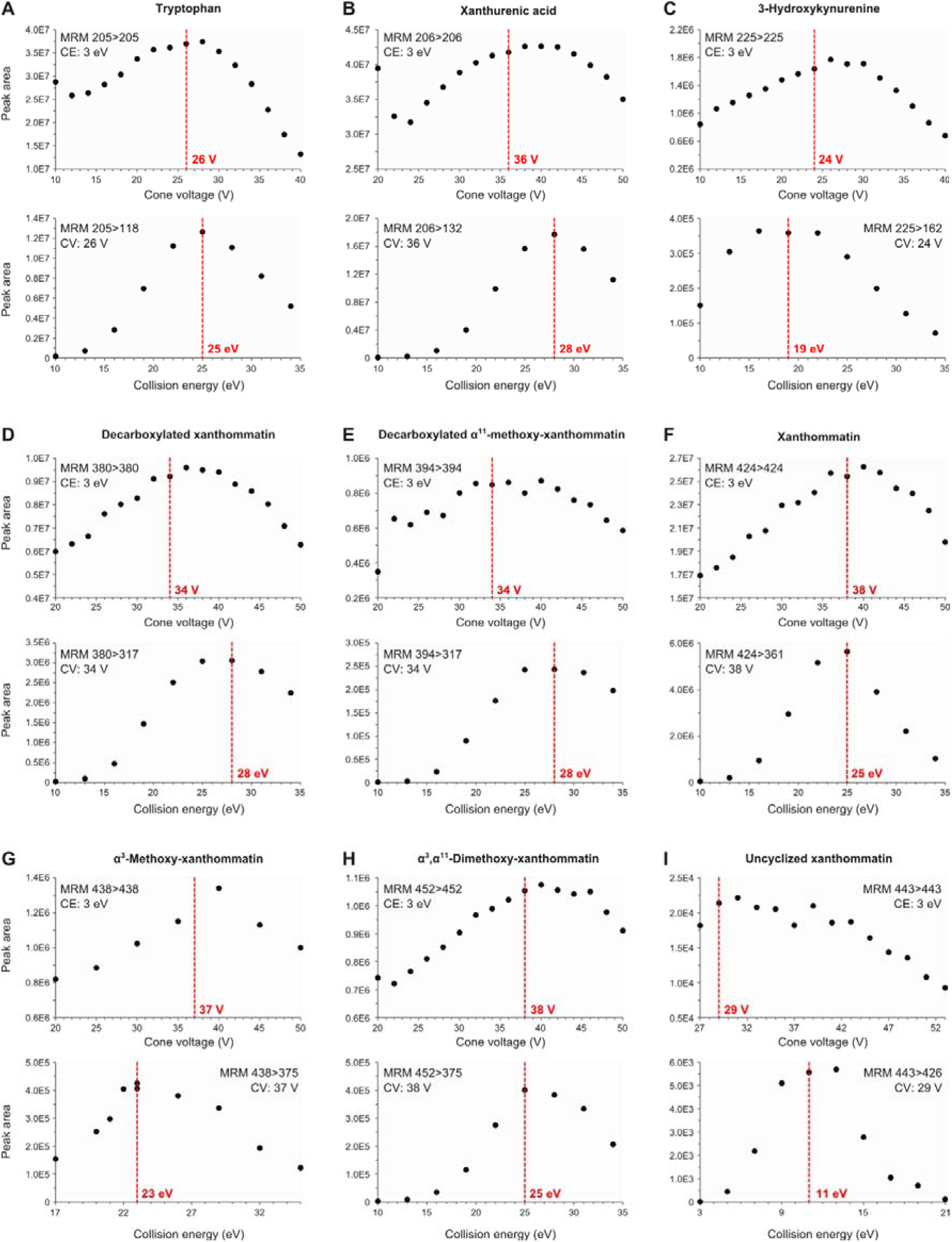
Optimization of cone voltages and collision energies for single reaction monitoring of ommochrome-related metabolites. Optimization of cone voltages (CV) was achieved by monitoring the parent-to-parent ion transition for different CV at the minimum collision energy (CE = 3 eV) during the same run. The optimum CV was defined as the CV just before the maximum peak area. Optimization of CE was achieved by monitoring a specific parent-to-product ion transition for different CE at the corresponding optimized CV during the same run. The optimum CE was defined as the CE corresponding to the maximum peak area.

**Fig S3.**
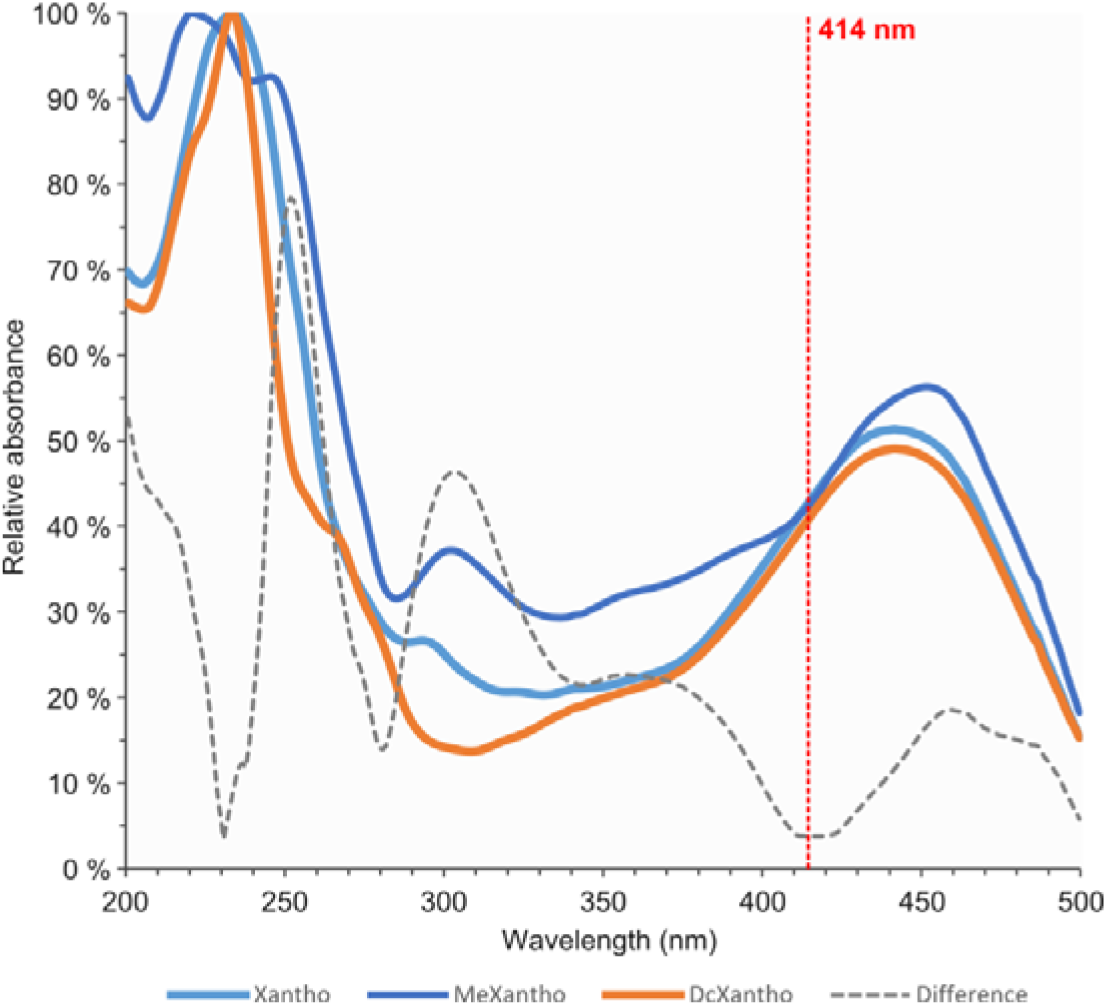
Determination of the wavelength at which ommatins absorb light equivalently. Synthesized ommatins incubated in acidified methanol were separated by liquid chromatography and analyzed by absorption spectroscopy. Relative absorbance spectra of xanthommatin (Xantho), α-methoxy-xanthommatin (MeXantho) and decarboxylated xanthommatin (DcXantho) were reported relatively to their respective maximum absorbance value. Gray dashed curve, difference between the three relative absorbance spectra calculated with the Manhattan formula: Difference(x) = |Abs_Xantho_(x) – Abs_MeXantho_(x)| + |Abs_Xantho_(x) – Abs_DcXantho_(x)| + |Abs_MeXantho_(x) – Abs_DcXantho_(x)|. The chromophores of the three ommatins absorb light equivalently at the wavelength for which this difference is minimal, i.e. 414 nm.

**Fig S4.**
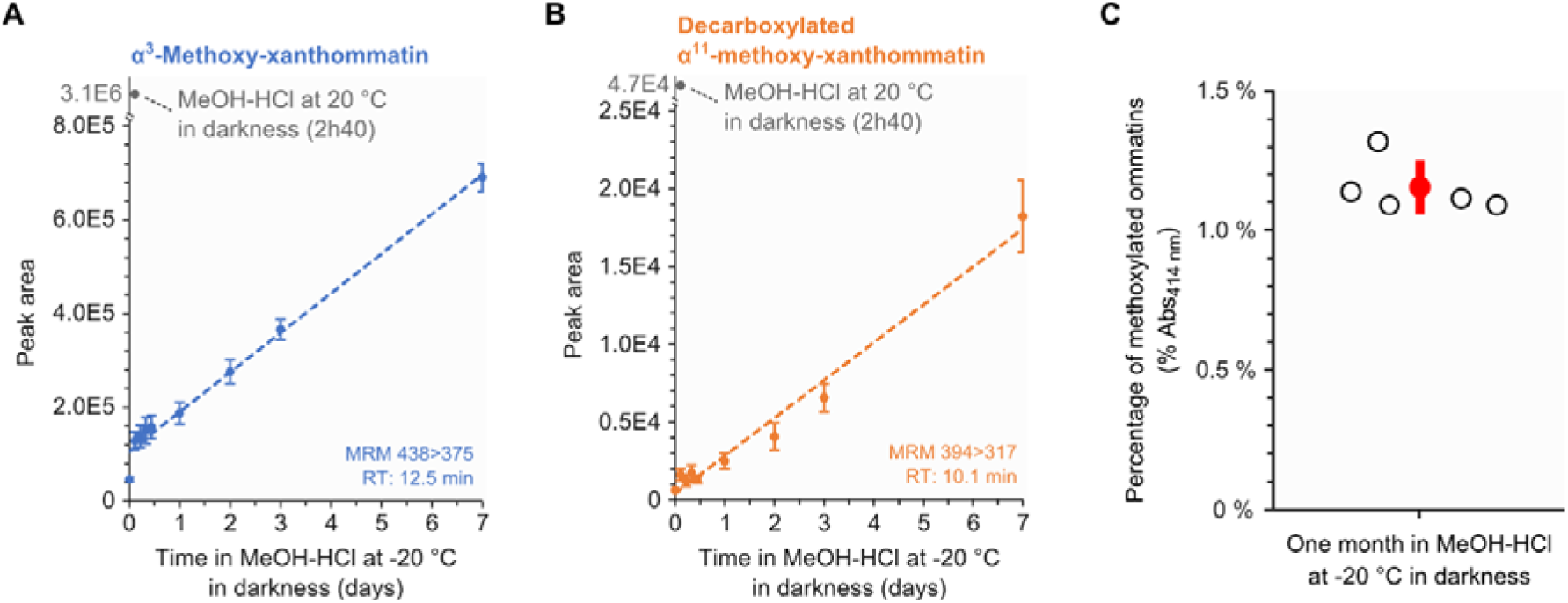
Slow methoxylations of synthesized ommatins in acidified methanol at −20 °C in darkness. Synthesized ommatins were solubilized in methanol acidified with 0.5 % (MeOH-HCl) and stored at −20 °C in darkness. Incubated ommatins were then separated by liquid chromatography. (**A**) Multiple reaction monitoring (MRM) signal of α-methoxy-xanthommatin. Values are mean ± S.D. of five samples. See File S2 for information on regression analyses. (**B**) MRM signal of decarboxylated α-methoxy-xanthommatin. Values are mean ± S.D. of five samples. See File S2 for information on regression analyses. **C)** Relative quantities of methoxylated ommatins (i.e. α-methoxy-xanthommatin, α, α-dimethoxy-xanthommatin and decarboxylated α-xanthommatin) in synthesized ommatins incubated in MeOH-HCl for one month at −20 °C in darkness. Quantifications were based on the absorbance of ommatins at 414 nm. Open circles, sample value. Red circle and error bar, mean value ± S.D. of five samples.

**Fig S5.**
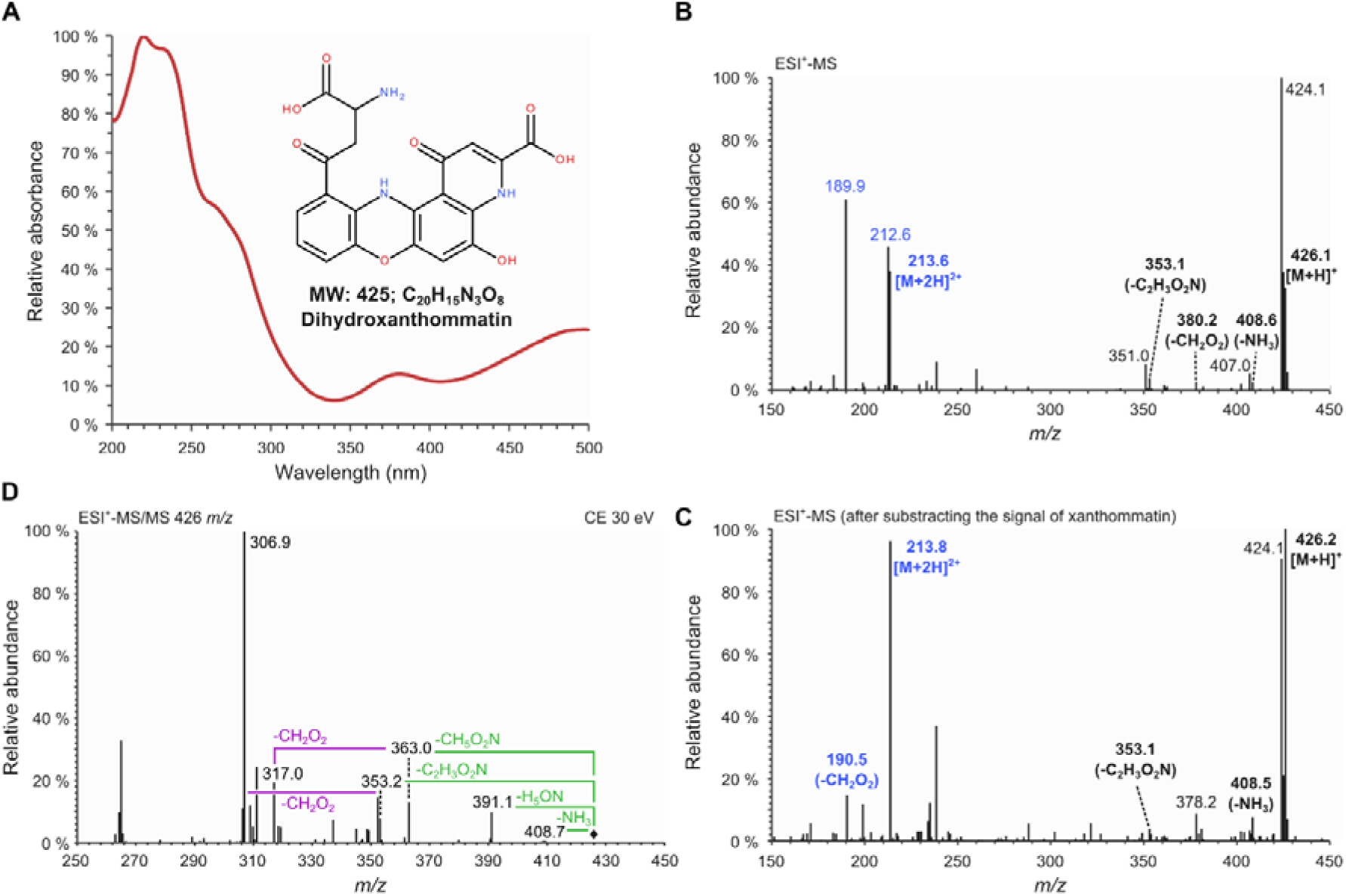
Analytical characterization of dihydroxanthommatin. Synthesized ommatins were solubilized in acidified methanol with 0.5 % HCl and separated by liquid chromatography (see File S1 for the experimental procedure). (**A**) Absorbance spectrum of dihydroxanthommatin. The peak at 480 nm makes dihydroxanthommatin appearing red in solution. (**B-C**) Raw mass spectrum (B) and mass spectrum with the signal of xanthommatin from the same run subtracted (C) of dihydroxanthommatin in positive mode. Ions associated to dihydroxanthommatin are indicated in bold fonts. Monocharged and double-charged ions are indicated in black and blue fonts, respectively. Mass losses of in-source fragments are indicated in parenthesis. (**D**) Tandem mass spectrum of dihydroxanthommatin obtained by the fragmentation of the molecular ion [M+H]^+^ 426 *m/z* at a collision energy (CE) of 30 eV. Losses corresponding to the typical fragmentation of the amino acid chain of ommatins are indicated in green. Losses corresponding to the fragmentation of the pyrido-carboxylic acid are reported in purple.

**Fig S6.**
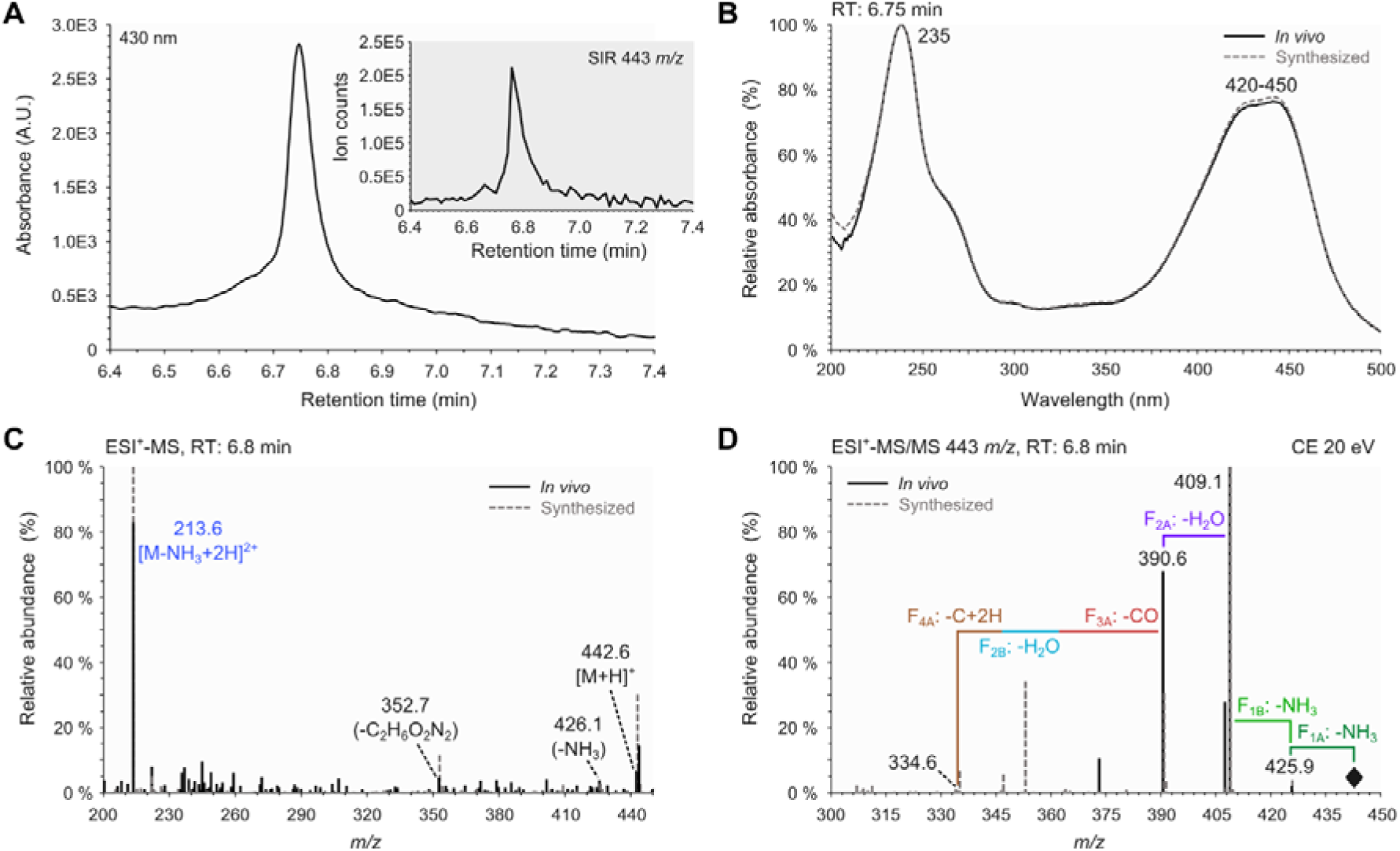
Analytical characterization of uncyclized xanthommatin in methanolic extracts of housefly eyes. (**A**) Chromatographic peak of absorbance at 430 nm. Inset, corresponding chromatographic peak of the associated [M+H]^+^ 443 *m/z* acquired in Single Ion Recording (SIR) mode. (**B**) Solid curve, absorbance spectrum corresponding to the chromatographic peak shown in panel A. Dashed curve, absorbance spectrum of synthesized uncyclized xanthommatin. (**C**) Solid lines, mass spectrum (MS) corresponding to the chromatographic peak shown in the inset of panel A. Dashed lines, MS of synthesized uncyclized xanthommatin. Black fonts, monocharged ions. Blue fonts, double-charged ion. (**D**) Solid lines, tandem mass (MS/MS) spectrum corresponding to the chromatographic peak shown in the inset of panel A. Dashed lines, MS/MS spectrum of synthesized uncyclized xanthommatin. F_1A_, F_1B_, F_2A_, F_2B_, F_3A_ and F_4A_ correspond to the fragmentations described in Fig 5.

**Fig S7.**
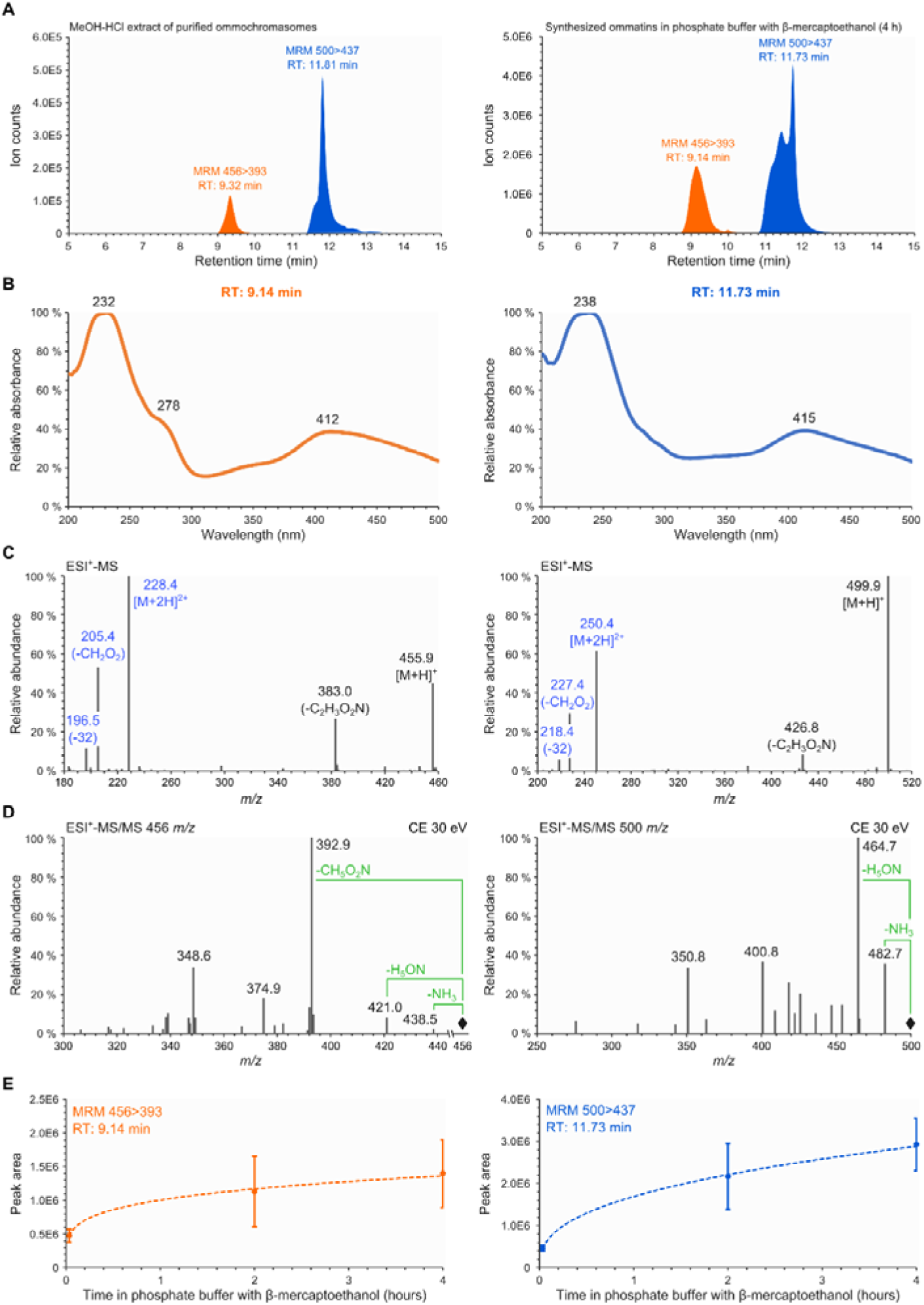
Ommatins are altered by addition of β-mercaptoethanol. Ommatins were separated by liquid chromatography and analysed by absorption, mass and tandem mass spectroscopies. (**A**) Chromatograms of acidified methanol (MeOH-HCl) extract of purified ommochromasomes and synthesized xanthommatin solubilized in phosphate buffer with β-mercaptoethanol. Only the signals of the multiple reaction monitoring (MRM) for the 456- and 500 *m/z*-associated compounds are shown. (**B-D**) Analytical characterization of the 456- and the 500-*m/z*-associated compounds present in phosphate buffer with β-mercaptoethanol. (B) Absorbance spectra. Wavelengths of peaks are reported. (C) Mass spectra in positive mode. Monocharged and double-charged ions are indicated in black and blue fonts, respectively. Mass losses of in-source fragments are indicated in parenthesis. (D) Tandem mass spectra in positive mode of the molecular ions [M+H]^+^. Main fragments are indicated in black fonts. Losses corresponding to the typical fragmentation of the amino acid chain of ommatins are reported in green. (**E**) Kinetics of formation of the 456- and 500-*m/z*-associated compounds in phosphate buffer with β-mercaptoethanol at 20 °C in darkness. The two compounds were monitored and quantified by MRM. Values are mean ± S.D. of four samples. See File S2 for information on regression analyses.

