## Supplemental File S1 for "Uncyclized xanthommatin is a key ommochrome intermediate in invertebrate coloration"

### Supplemental File S1 – Supplemental Material and Methods

#### Analytical characterization of dihydroxanthommatin

Dihydroxanthommatin, the reduced form of xanthommatin, was not detectable using the CSH™ C18 column, most likely because dihydroxanthommatin was oxidized into xanthommatin by the chemistry of the column. Therefore, we analytically characterized dihydroxanthommatin after separation on a HSS T3 C18 column (2.1 x 150 mm, 1.7 µm) equipped with a HSS T3 C18 VanGuard™ pre-column (2.1 x 5 mm). All other analytical conditions were the same than those described in the experimental procedure section.

#### Reactivity of synthesized ommatins with β-mercaptoethanol

##### *Sample preparation*

Solutions of synthesized ommatins were prepared at 1 mg/mL in phosphate buffer 0.2 M, pH 7.0 and 14 mM β-mercaptoethanol (n = 4). The samples were mixed for 30 s and filtered using 0.45 µm filters. Aliquots of 50 µL were prepared for each sample and stored in darkness at 20 °C. Each aliquot was analyzed only once and represented a single time point for each sample.

##### *Quantifications*

β-Mercaptoethanol-added forms were detected and quantified by MS/MS (multiple reaction monitoring [MRM] mode) spectrometry. The following parent-to-product ion transitions were monitored: $[M+H]^+$ 456>393 *m/z* (cone voltage 43 V, collision energy 25 eV) and $[M+H]^+$ 500>437 *m/z* (cone voltage 43 V, collision energy 25 eV). Peak areas were calculated by integrating chromatographic peaks with a “Mean” smoothing method (window size: ± 3 scans, number of smooths: 2).
