## Supplemental File S2 for "Uncyclized xanthommatin is a key ommochrome intermediate in invertebrate coloration"

### Supplemental File S2 – Statistical analyses

##### Statistics of Figure 4A-E

```
mrm = read.table("MRM_MeOH-HCl_20_RData.csv", h=T, sep=";")

### Kinetic of xanthommatin
cor.test(mrm$Time_h, mrm$MRM_424, method = "spearman")

## Warning in cor.test.default(mrm$Time_h, mrm$MRM_424, method = "spearman"
):
## Cannot compute exact p-value with ties

##
##  Spearman's rank correlation rho
##
## data:  mrm$Time_h and mrm$MRM_424
## S = 12742, p-value < 2.2e-16
## alternative hypothesis: true rho is not equal to 0
## sample estimates:
##        rho
## -0.9468069

plot(mrm$MRM_424~mrm$Time_h)
abline(lm(mrm$MRM_424~mrm$Time_h))
```

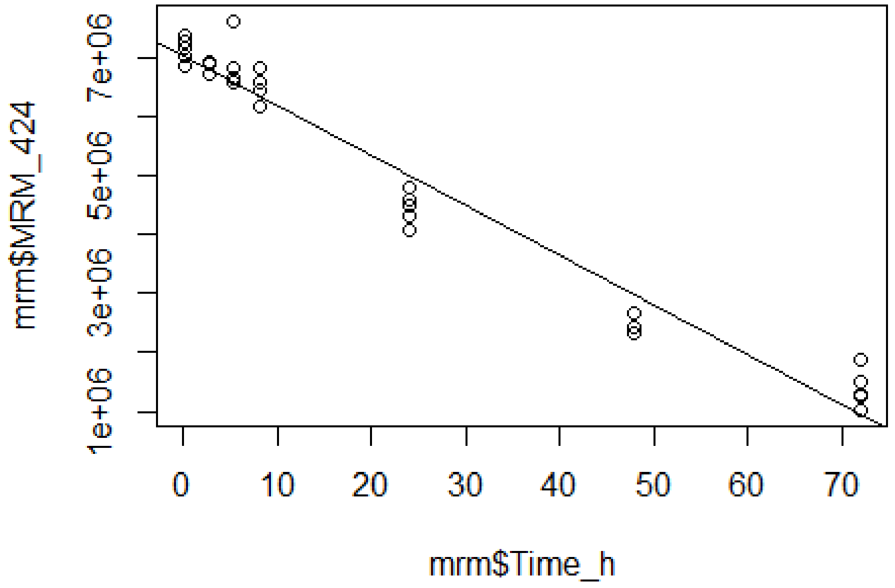


```
summary(lm(mrm$MRM_424~mrm$Time_h))

##
## Call:
## lm(formula = mrm$MRM_424 ~ mrm$Time_h)
##
```

```
## Residuals:
##     Min      1Q  Median      3Q     Max 
## -926615 -193228   38852  248987 1049135 
## 
## Coefficients:
##             Estimate Std. Error t value Pr(>|t|)    
## (Intercept)  7029781      99448   70.69   <2e-16 ***
## mrm$Time_h    -84278       2955  -28.52   <2e-16 ***
## ---
## Signif. codes:  0 '***' 0.001 '**' 0.01 '*' 0.05 '.' 0.1 ' ' 1
## 
## Residual standard error: 437000 on 32 degrees of freedom
##   (1 observation deleted due to missingness)
## Multiple R-squared:  0.9622, Adjusted R-squared:  0.961 
## F-statistic: 813.5 on 1 and 32 DF,  p-value: < 2.2e-16



### Kinetic of methoxy-xanthommatin
cor.test(mrm$Time_h, mrm$MRM_438, method = "spearman")

## Warning in cor.test.default(mrm$Time_h, mrm$MRM_438, method = "spearman"
): 
## Cannot compute exact p-value with ties

## 
##  Spearman's rank correlation rho
## 
## data:  mrm$Time_h and mrm$MRM_438
## S = 119.87, p-value < 2.2e-16
## alternative hypothesis: true rho is not equal to 0
## sample estimates:
##       rho 
## 0.9816852

plot(mrm$MRM_438~log(mrm$Time_h))
abline(lm(mrm$MRM_438~log(mrm$Time_h)))
```

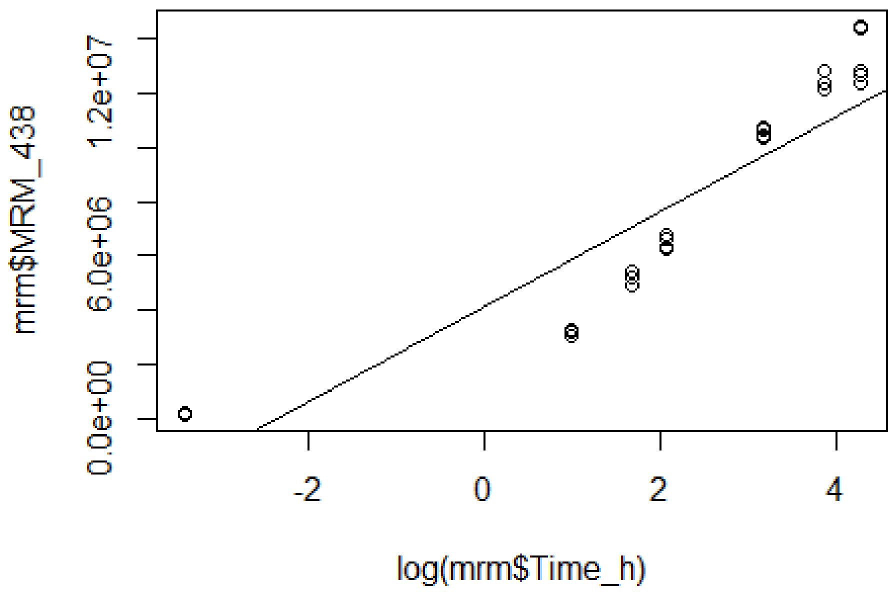


```
summary(lm(mrm$MRM_438~log(mrm$Time_h)))

##
## Call:
## lm(formula = mrm$MRM_438 ~ log(mrm$Time_h))
##
## Residuals:
##      Min       1Q   Median       3Q      Max
## -2812242 -1780376   757895  1830503  2881893
##
## Coefficients:
##                    Estimate Std. Error t value Pr(>|t|)
## (Intercept)         4114164     410369   10.03 2.13e-11 ***
## log(mrm$Time_h)     1744666     138031   12.64 5.51e-14 ***
## ---
## Signif. codes:  0 '***' 0.001 '**' 0.01 '*' 0.05 '.' 0.1 ' ' 1
##
## Residual standard error: 1937000 on 32 degrees of freedom
##   (1 observation deleted due to missingness)
## Multiple R-squared:  0.8331, Adjusted R-squared:  0.8279
## F-statistic: 159.8 on 1 and 32 DF,  p-value: 5.512e-14



### Kinetic of dimethoxy-xanthommatin
cor.test(mrm$Time_h, mrm$MRM_452, method = "spearman")

## Warning in cor.test.default(mrm$Time_h, mrm$MRM_452, method = "spearman"
## ):
## Cannot compute exact p-value with ties
```

```
##
##  Spearman's rank correlation rho
##
## data:  mrm$Time_h and mrm$MRM_452
## S = 55.378, p-value < 2.2e-16
## alternative hypothesis: true rho is not equal to 0
## sample estimates:
##       rho
## 0.9863602

plot(mrm$MRM_452~mrm$Time_h)
abline(lm(mrm$MRM_452~mrm$Time_h))
```

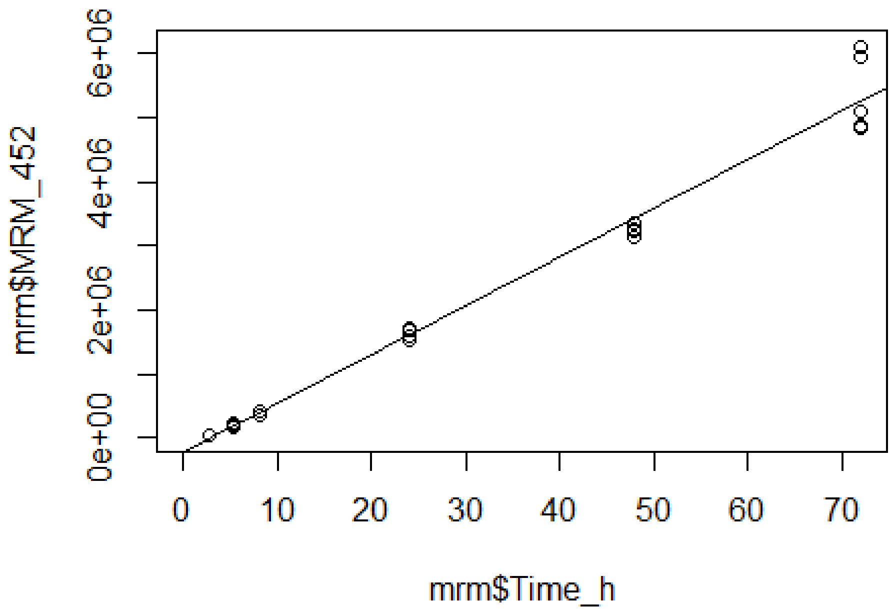


```
summary(lm(mrm$MRM_452~mrm$Time_h))

##
## Call:
## lm(formula = mrm$MRM_452 ~ mrm$Time_h)
##
## Residuals:
##     Min      1Q  Median      3Q     Max
## -440670  -84604  -11250   56433  838647
##
## Coefficients:
##              Estimate Std. Error t value Pr(>|t|)
## (Intercept)  -207533      67957  -3.054  0.00503 **
## mrm$Time_h     76101       1865  40.810  < 2e-16 ***
## ---
## Signif. codes:  0 '***' 0.001 '**' 0.01 '*' 0.05 '.' 0.1 ' ' 1
##
## Residual standard error: 257100 on 27 degrees of freedom
```

```
##   (6 observations deleted due to missingness)
## Multiple R-squared:  0.984,  Adjusted R-squared:  0.9835
## F-statistic:  1665 on 1 and 27 DF,  p-value: < 2.2e-16



### Kinetic of decarboxylated xanthommatin
cor.test(mrm$Time_h, mrm$MRM_380, method = "spearman")

## Warning in cor.test.default(mrm$Time_h, mrm$MRM_380, method = "spearman"
): 
## Cannot compute exact p-value with ties

## 
##  Spearman's rank correlation rho
## 
## data:  mrm$Time_h and mrm$MRM_380
## S = 2415.8, p-value = 6.333e-05
## alternative hypothesis: true rho is not equal to 0
## sample estimates:
##       rho 
## 0.6308959

plot(mrm$MRM_380~log(mrm$Time_h))
abline(lm(mrm$MRM_380~log(mrm$Time_h)))
```

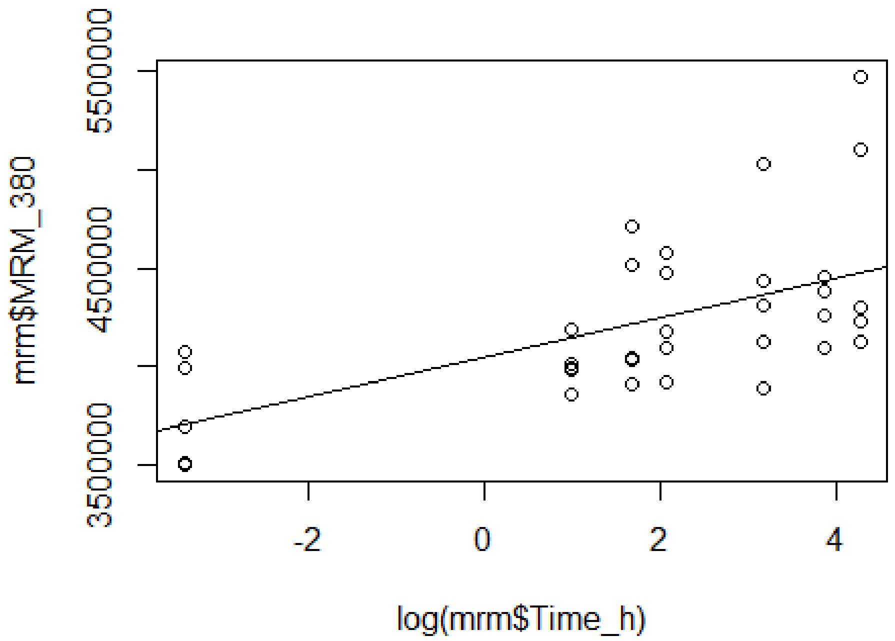


```
summary(lm(mrm$MRM_380~log(mrm$Time_h)))

## 
## Call:
## lm(formula = mrm$MRM_380 ~ log(mrm$Time_h))
## 
```

```
## Residuals:
##     Min      1Q  Median      3Q     Max
## -475442 -197184 -138365  186024  997024
##
## Coefficients:
##                   Estimate Std. Error t value Pr(>|t|)
## (Intercept)        4044202      72209  56.007  < 2e-16 ***
## log(mrm$Time_h)     100599      24288   4.142 0.000235 ***
## ---
## Signif. codes:  0 '***' 0.001 '**' 0.01 '*' 0.05 '.' 0.1 ' ' 1
##
## Residual standard error: 340800 on 32 degrees of freedom
##   (1 observation deleted due to missingness)
## Multiple R-squared:  0.349,  Adjusted R-squared:  0.3287
## F-statistic: 17.16 on 1 and 32 DF,  p-value: 0.0002347



### Kinetic of decarboxylated methoxy-xanthommatin
cor.test(mrm$Time_h, mrm$MRM_394, method = "spearman")

## Warning in cor.test.default(mrm$Time_h, mrm$MRM_394, method = "spearman"
## ):
## Cannot compute exact p-value with ties

##
##  Spearman's rank correlation rho
##
## data:  mrm$Time_h and mrm$MRM_394
## S = 85.528, p-value < 2.2e-16
## alternative hypothesis: true rho is not equal to 0
## sample estimates:
##       rho
## 0.9869324

plot(mrm$MRM_394~mrm$Time_h)
abline(lm(mrm$MRM_394~mrm$Time_h))
```

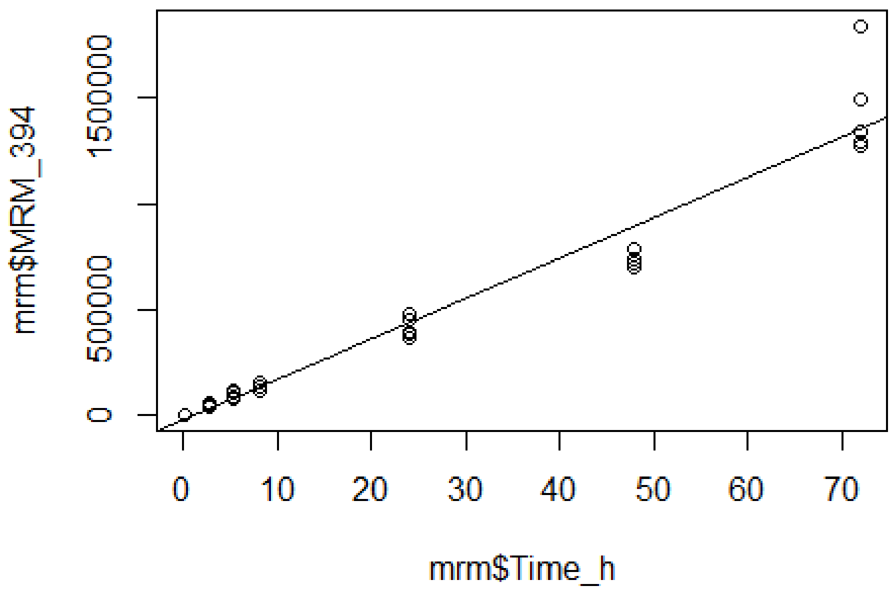


```
summary(lm(mrm$MRM_394~mrm$Time_h))

##
## Call:
## lm(formula = mrm$MRM_394 ~ mrm$Time_h)
##
## Residuals:
##     Min      1Q  Median      3Q     Max
## -200036  -33979   11410   26021  484111
##
## Coefficients:
##              Estimate Std. Error t value Pr(>|t|)
## (Intercept) -24727.5    25291.7  -0.978    0.336
## mrm$Time_h   19176.2      751.5  25.518   <2e-16 ***
## ---
## Signif. codes:  0 '***' 0.001 '**' 0.01 '*' 0.05 '.' 0.1 ' ' 1
##
## Residual standard error: 111100 on 32 degrees of freedom
##   (1 observation deleted due to missingness)
## Multiple R-squared:  0.9532, Adjusted R-squared:  0.9517
## F-statistic: 651.2 on 1 and 32 DF,  p-value: < 2.2e-16
```

**Statistics of Figure 4F**

```
abs = read.table("414nm_RData.csv", h=T, sep=";")
abs_2 = subset(abs, abs$Time==2)
abs_480 = subset(abs, abs$Time==480)
abs_1440 = subset(abs, abs$Time==1440)
abs_4320 = subset(abs, abs$Time==4320)


### Normality analysis
shapiro.test(abs_2$Unaltered_ommatins) # NO

##
##  Shapiro-Wilk normality test
##
## data:  abs_2$Unaltered_ommatins
## W = 0.63564, p-value = 0.00179

shapiro.test(abs_480$Unaltered_ommatins) # OK

##
##  Shapiro-Wilk normality test
##
## data:  abs_480$Unaltered_ommatins
## W = 0.79418, p-value = 0.0726

shapiro.test(abs_1440$Unaltered_ommatins) # NO

##
##  Shapiro-Wilk normality test
##
## data:  abs_1440$Unaltered_ommatins
## W = 0.61851, p-value = 0.001093

shapiro.test(abs_4320$Unaltered_ommatins) # NO

##
##  Shapiro-Wilk normality test
##
## data:  abs_4320$Unaltered_ommatins
## W = 0.65141, p-value = 0.002767



### Homoscedasticity analysis
fligner.test(formula(abs$Unaltered_ommatins~abs$Time)) # OK

##
##  Fligner-Killeen test of homogeneity of variances
##
## data:  abs$Unaltered_ommatins by abs$Time
## Fligner-Killeen:med chi-squared = 4.5139, df = 3, p-value = 0.2111



### Median analysis
kruskal.test(formula(abs$Unaltered_ommatins~abs$Time))
```

```
##
##  Kruskal-Wallis rank sum test
##
## data:  abs$Unaltered_ommatins by abs$Time
## Kruskal-Wallis chi-squared = 17.857, df = 3, p-value = 0.0004707

pairwise.wilcox.test(abs$Unaltered_ommatins, abs$Time)

##
##  Pairwise comparisons using Wilcoxon rank sum test
##
## data:  abs$Unaltered_ommatins and abs$Time
##
##      2     480   1440
## 480  0.048 -     -
## 1440 0.048 0.048 -
## 4320 0.048 0.048 0.048
##
## P value adjustment method: holm
```

**Statistics of Figure 5A-B**

```
ux = read.table("UncXantho_kinetic_RData.csv", h=T, sep=";")


### Kinetic of DAD signal
cor.test(ux$Time, ux$DAD, method="spearman")

## Warning in cor.test.default(ux$Time, ux$DAD, method = "spearman"): Canno
t
## compute exact p-value with ties

##
##  Spearman's rank correlation rho
##
## data:  ux$Time and ux$DAD
## S = 27353, p-value = 3.587e-11
## alternative hypothesis: true rho is not equal to 0
## sample estimates:
##        rho
## -0.8018962

plot(ux$DAD~log(ux$Time))
abline(lm(ux$DAD~log(ux$Time)))
```

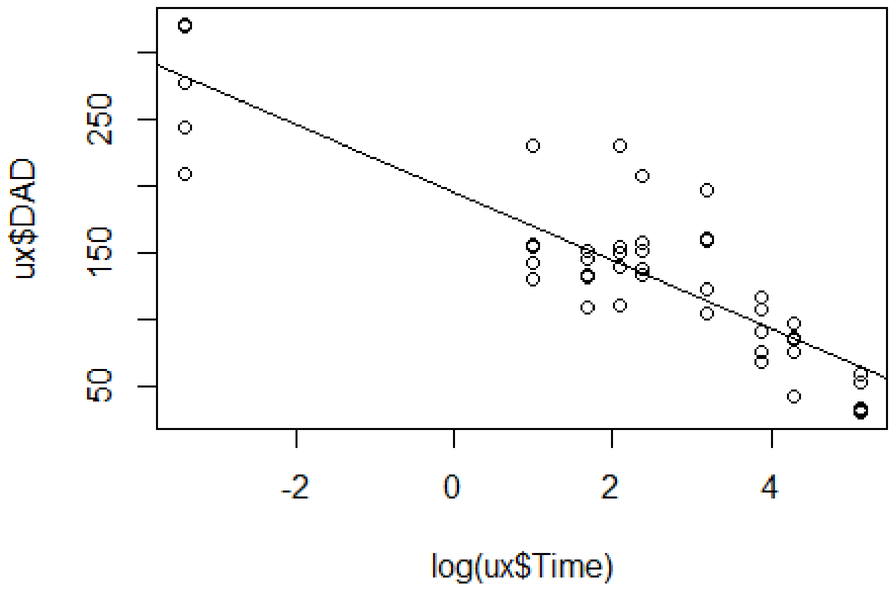


```
summary(lm(ux$DAD~log(ux$Time)))

##
## Call:
## lm(formula = ux$DAD ~ log(ux$Time))
##
## Residuals:
```

```
##      Min       1Q  Median      3Q     Max
## -73.098 -21.649  -5.098  12.723  88.723
##
## Coefficients:
##               Estimate Std. Error t value Pr(>|t|)
## (Intercept)    195.230      7.215   27.06  < 2e-16 ***
## log(ux$Time)   -25.465      2.221  -11.46 1.16e-14 ***
## ---
## Signif. codes:  0 '***' 0.001 '**' 0.01 '*' 0.05 '.' 0.1 ' ' 1
##
## Residual standard error: 35.08 on 43 degrees of freedom
## Multiple R-squared:  0.7535, Adjusted R-squared:  0.7478
## F-statistic: 131.4 on 1 and 43 DF,  p-value: 1.163e-14
```

```
### Kinetic of MRM signal
cor.test(ux$Time, ux$SIR, method="spearman")
```

```
## Warning in cor.test.default(ux$Time, ux$SIR, method = "spearman"): Canno
t
## compute exact p-value with ties
```

```
##
##  Spearman's rank correlation rho
##
## data:  ux$Time and ux$SIR
## S = 27564, p-value = 8.723e-12
## alternative hypothesis: true rho is not equal to 0
## sample estimates:
##        rho
## -0.8157865
```

```
plot(ux$SIR~log(ux$Time))
abline(lm(ux$SIR~log(ux$Time)))
```

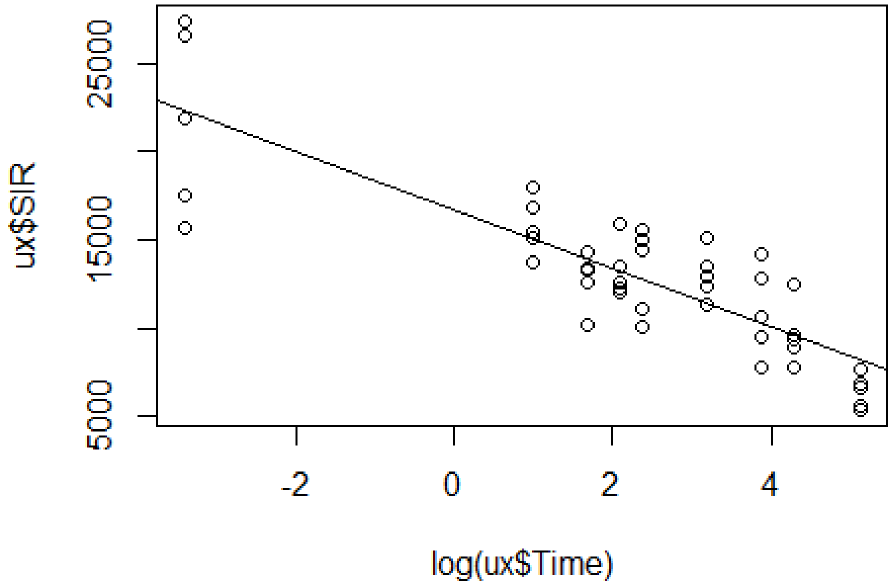


```
summary(lm(ux$SIR~log(ux$Time)))

##
## Call:
## lm(formula = ux$SIR ~ log(ux$Time))
##
## Residuals:
##     Min      1Q  Median      3Q     Max
## -6626.3 -1340.2  -255.8  1795.8  5079.7
##
## Coefficients:
##              Estimate Std. Error t value Pr(>|t|)
## (Intercept)   16674.9      506.0   32.95  < 2e-16 ***
## log(ux$Time)  -1645.0      155.8  -10.56 1.61e-13 ***
## ---
## Signif. codes:  0 '***' 0.001 '**' 0.01 '*' 0.05 '.' 0.1 ' ' 1
##
## Residual standard error: 2460 on 43 degrees of freedom
## Multiple R-squared:  0.7217, Adjusted R-squared:  0.7152
## F-statistic: 111.5 on 1 and 43 DF,  p-value: 1.61e-13
```

**Statistics of Figure 6C, E**

```
ratios = read.table("RatiosOHK_RData.csv", h=T, sep=";")

OHKtoXA = subset(ratios, ratios$Ratio=="OHKtoXA")
OHKtoUX = subset(ratios, ratios$Ratio=="OHKtoUncX")

FlyHeads = subset(ratios, ratios$Category=="FlyHeads")
Ommochromasomes = subset(ratios, ratios$Category=="Ommochromasomes")

### Normality analysis
shapiro.test(subset(OHKtoXA, OHKtoXA$Category=="FlyHeads")$Value) # OK

##
##  Shapiro-Wilk normality test
##
## data:  subset(OHKtoXA, OHKtoXA$Category == "FlyHeads")$Value
## W = 0.84019, p-value = 0.1655

shapiro.test(subset(OHKtoXA, OHKtoXA$Category=="Ommochromasomes")$Value) #
OK

##
##  Shapiro-Wilk normality test
##
## data:  subset(OHKtoXA, OHKtoXA$Category == "Ommochromasomes")$Value
## W = 0.89859, p-value = 0.4022

shapiro.test(subset(OHKtoUX, OHKtoUX$Category=="FlyHeads")$Value) # OK

##
##  Shapiro-Wilk normality test
##
## data:  subset(OHKtoUX, OHKtoUX$Category == "FlyHeads")$Value
## W = 0.85101, p-value = 0.1977

shapiro.test(subset(OHKtoUX, OHKtoUX$Category=="Ommochromasomes")$Value) #
OK

##
##  Shapiro-Wilk normality test
##
## data:  subset(OHKtoUX, OHKtoUX$Category == "Ommochromasomes")$Value
## W = 0.95313, p-value = 0.7595

### Homoscedasticity analysis
fligner.test(formula(OHKtoXA$Value~OHKtoXA$Category)) # OK

##
##  Fligner-Killeen test of homogeneity of variances
##
## data:  OHKtoXA$Value by OHKtoXA$Category
## Fligner-Killeen:med chi-squared = 0.0039292, df = 1, p-value = 0.95

fligner.test(formula(OHKtoUX$Value~OHKtoUX$Category)) # OK
```

```
##
##  Fligner-Killeen test of homogeneity of variances
##
## data:  OHKtoUX$Value by OHKtoUX$Category
## Fligner-Killeen:med chi-squared = 3.194, df = 1, p-value = 0.07391
```

```
fligner.test(formula(FlyHeads$Value~FlyHeads$Ratio)) # OK
```

```
##
##  Fligner-Killeen test of homogeneity of variances
##
## data:  FlyHeads$Value by FlyHeads$Ratio
## Fligner-Killeen:med chi-squared = 1.9797, df = 1, p-value = 0.1594
```

```
fligner.test(formula(Ommochromasomes$Value~Ommochromasomes$Ratio)) # OK
```

```
##
##  Fligner-Killeen test of homogeneity of variances
##
## data:  Ommochromasomes$Value by Ommochromasomes$Ratio
## Fligner-Killeen:med chi-squared = 1.9797, df = 1, p-value = 0.1594
```

```
### Mean analysis
## Ratios of xanthurenic acid between crude extracts and ommochromasomes
t.test(subset(OHKtoXA, OHKtoXA$Category=="FlyHeads")$Value, subset(OHKtoXA,
OHKtoXA$Category=="Ommochromasomes")$Value, paired=TRUE)
```

```
##
##  Paired t-test
##
## data:  subset(OHKtoXA, OHKtoXA$Category == "FlyHeads")$Value and subset(
OHKtoXA, OHKtoXA$Category == "Ommochromasomes")$Value
## t = -8.2177, df = 4, p-value = 0.001195
## alternative hypothesis: true difference in means is not equal to 0
## 95 percent confidence interval:
##  -12.007645  -5.942842
## sample estimates:
## mean of the differences
##               -8.975243
```

```
## Ratios of uncyclized xanthommatin between crude extracts and ommochromas
omes
t.test(subset(OHKtoUX, OHKtoUX$Category=="FlyHeads")$Value, subset(OHKtoUX,
OHKtoUX$Category=="Ommochromasomes")$Value, paired=TRUE)
```

```
##
##  Paired t-test
##
## data:  subset(OHKtoUX, OHKtoUX$Category == "FlyHeads")$Value and subset(
OHKtoUX, OHKtoUX$Category == "Ommochromasomes")$Value
## t = 6.0573, df = 4, p-value = 0.003749
## alternative hypothesis: true difference in means is not equal to 0
## 95 percent confidence interval:
##   7.763434 20.903115
## sample estimates:
```

```
## mean of the differences
##                14.33327

## Ratios of xanturenic acid compared to uncyclized xanthommatin in crude e
xtracts
t.test(subset(FlyHeads, FlyHeads$Ratio=="OHKtoXA")$Value, subset(Ommochroma
somes, Ommochromasomes$Ratio=="OHKtoXA")$Value, var.equal = TRUE)

##
##  Two Sample t-test
##
## data:  subset(FlyHeads, FlyHeads$Ratio == "OHKtoXA")$Value and subset(Om
mochromasomes, Ommochromasomes$Ratio == "OHKtoXA")$Value
## t = -8.8831, df = 8, p-value = 2.04e-05
## alternative hypothesis: true difference in means is not equal to 0
## 95 percent confidence interval:
##  -11.305159  -6.645328
## sample estimates:
## mean of x mean of y
##  2.083669 11.058912

## Ratios of xanturenic acid compared to uncyclized xanthommatin in ommochr
omasomes
t.test(subset(FlyHeads, FlyHeads$Ratio=="OHKtoUncX")$Value, subset(Ommochro
masomes, Ommochromasomes$Ratio=="OHKtoUncX")$Value, var.equal = TRUE)

##
##  Two Sample t-test
##
## data:  subset(FlyHeads, FlyHeads$Ratio == "OHKtoUncX")$Value and subset(
Ommochromasomes, Ommochromasomes$Ratio == "OHKtoUncX")$Value
## t = 5.7091, df = 8, p-value = 0.0004496
## alternative hypothesis: true difference in means is not equal to 0
## 95 percent confidence interval:
##   8.543833 20.122716
## sample estimates:
## mean of x mean of y
## 20.516923  6.183649
```

**Statistics on proportions of decarboxylated xanthommatin**

```r
abs = read.table("414nm_DcOmmatins_RData.csv", h=T, sep=";")

invitro = subset(abs, abs$Type=="InVitro")
omm = subset(abs, abs$Type=="Ommochromasome")


### Normality analysis
shapiro.test(invitro$pDcOmmatins) # OK

##
##  Shapiro-Wilk normality test
##
## data:  invitro$pDcOmmatins
## W = 0.97263, p-value = 0.9141

shapiro.test(omm$pDcOmmatins) # OK

##
##  Shapiro-Wilk normality test
##
## data:  omm$pDcOmmatins
## W = 0.83902, p-value = 0.1622

### Homoscedasticity analysis
fligner.test(formula(abs$pDcOmmatins~abs$Type)) # NO

##
##  Fligner-Killeen test of homogeneity of variances
##
## data:  abs$pDcOmmatins by abs$Type
## Fligner-Killeen:med chi-squared = 4.2109, df = 1, p-value = 0.04017

### Mean analysis
t.test(invitro$pDcOmmatins, omm$pDcOmmatins, var.equal = FALSE)

##
##  Welch Two Sample t-test
##
## data:  invitro$pDcOmmatins and omm$pDcOmmatins
## t = 219.99, df = 12.499, p-value < 2.2e-16
## alternative hypothesis: true difference in means is not equal to 0
## 95 percent confidence interval:
##  0.1602019 0.1633927
## sample estimates:
## mean of x mean of y
## 0.2152880 0.0534907
```

**Statistics of Figure S4A-B**

```
mrm = read.table("MRM_MeOH-HCl_-20_RData.csv", h=T, sep=";")


### Kinetic of decarboxylated methoxy-xanthommatin
cor.test(mrm$Time_h, mrm$MRM_394, method = "spearman")

## Warning in cor.test.default(mrm$Time_h, mrm$MRM_394, method = "spearman"
):
## Cannot compute exact p-value with ties

##
##  Spearman's rank correlation rho
##
## data:  mrm$Time_h and mrm$MRM_394
## S = 2414.1, p-value = 4.834e-13
## alternative hypothesis: true rho is not equal to 0
## sample estimates:
##       rho
## 0.8409692

plot(mrm$MRM_394~mrm$Time_h)
abline(lm(mrm$MRM_394~mrm$Time_h))
```

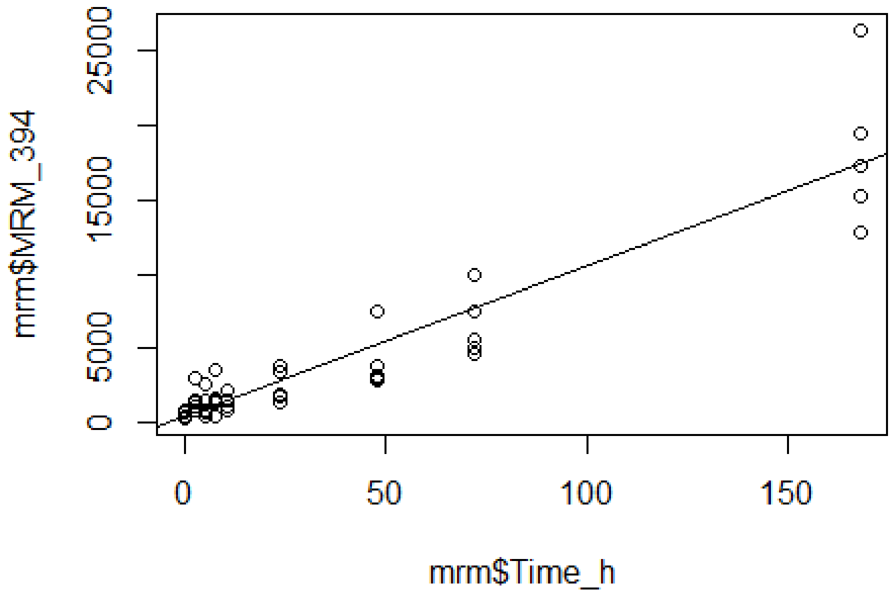


```
summary(lm(mrm$MRM_394~mrm$Time_h))

##
## Call:
## lm(formula = mrm$MRM_394 ~ mrm$Time_h)
```

```
##
## Residuals:
##     Min      1Q  Median      3Q     Max
## -4618.5  -949.2    27.8   640.8  8985.5
##
## Coefficients:
##             Estimate Std. Error t value Pr(>|t|)
## (Intercept)  409.355    380.730   1.075    0.288
## mrm$Time_h   101.203      5.978  16.928   <2e-16 ***
## ---
## Signif. codes:  0 '***' 0.001 '**' 0.01 '*' 0.05 '.' 0.1 ' ' 1
##
## Residual standard error: 2060 on 43 degrees of freedom
## Multiple R-squared:  0.8695, Adjusted R-squared:  0.8665
## F-statistic: 286.6 on 1 and 43 DF,  p-value: < 2.2e-16
```

```
### Kinetic of methoxy-xanthommatin
cor.test(mrm$Time_h, mrm$MRM_438, method = "spearman")
```

```
## Warning in cor.test.default(mrm$Time_h, mrm$MRM_438, method = "spearman"
):
## Cannot compute exact p-value with ties
```

```
##
##  Spearman's rank correlation rho
##
## data:  mrm$Time_h and mrm$MRM_438
## S = 1770.3, p-value = 9.56e-16
## alternative hypothesis: true rho is not equal to 0
## sample estimates:
##       rho
## 0.8833822
```

```
plot(mrm$MRM_438~mrm$Time_h)
abline(lm(mrm$MRM_438~mrm$Time_h))
```

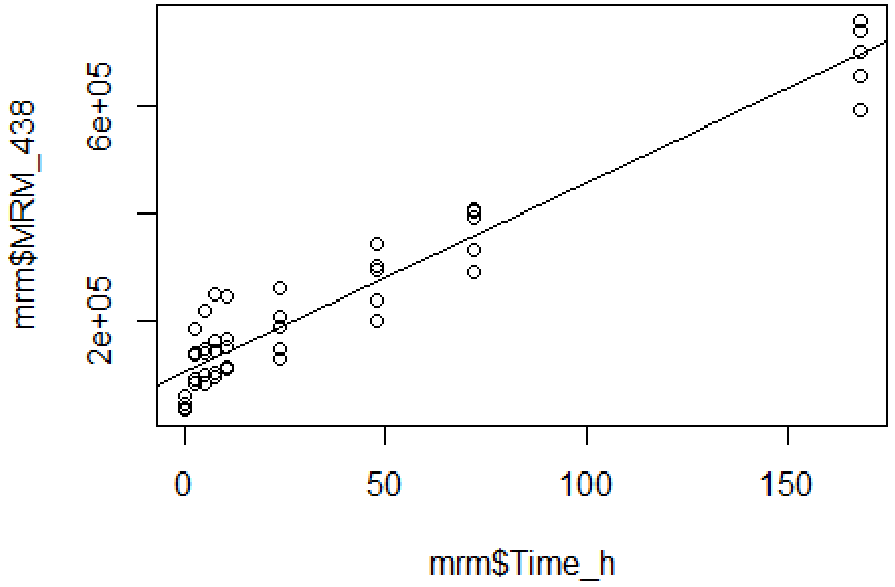


```
summary(lm(mrm$MRM_438~mrm$Time_h))

##
## Call:
## lm(formula = mrm$MRM_438 ~ mrm$Time_h)
##
## Residuals:
##     Min      1Q  Median      3Q     Max
## -104717  -38792    7543   28418  117089
##
## Coefficients:
##             Estimate Std. Error t value Pr(>|t|)
## (Intercept) 105925.5     9821.8   10.79  8.3e-14 ***
## mrm$Time_h    3514.9      154.2   22.79  < 2e-16 ***
## ---
## Signif. codes:  0 '***' 0.001 '**' 0.01 '*' 0.05 '.' 0.1 ' ' 1
##
## Residual standard error: 53150 on 43 degrees of freedom
## Multiple R-squared:  0.9235, Adjusted R-squared:  0.9218
## F-statistic: 519.4 on 1 and 43 DF,  p-value: < 2.2e-16
```

**Statistics of Figure S7E**

```
mrm = read.table("MRM_BME_RData.csv", h=T, sep=";")


### Kinetic of 456-m/z-associated compound
cor.test(mrm$Time_h, mrm$MRM_456, method = "spearman")

## Warning in cor.test.default(mrm$Time_h, mrm$MRM_456, method = "spearman"
## ):
## Cannot compute exact p-value with ties

##
##  Spearman's rank correlation rho
##
## data:  mrm$Time_h and mrm$MRM_456
## S = 91.517, p-value = 0.01497
## alternative hypothesis: true rho is not equal to 0
## sample estimates:
##       rho
## 0.6800093

plot(mrm$MRM_456~log(mrm$Time_h))
abline(lm(mrm$MRM_456~log(mrm$Time_h)))
```

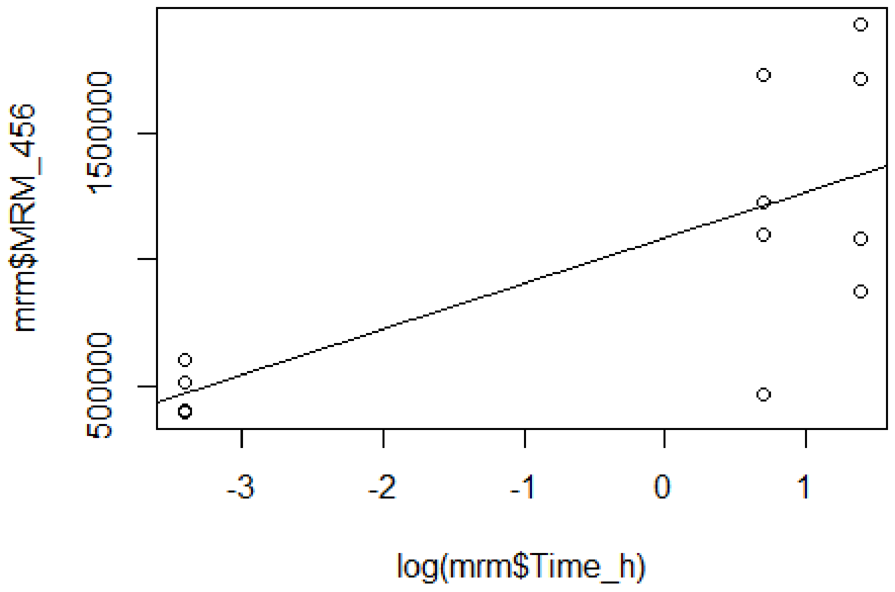


```
summary(lm(mrm$MRM_456~log(mrm$Time_h)))

##
## Call:
## lm(formula = mrm$MRM_456 ~ log(mrm$Time_h))
##
## Residuals:
```

```
##      Min       1Q  Median       3Q      Max
## -744762 -146675  -21520  196078  600035
##
## Coefficients:
##                 Estimate Std. Error t value Pr(>|t|)
## (Intercept)      1083850     120119   9.023 4.04e-06 ***
## log(mrm$Time_h)   181344      55528   3.266  0.00849 **
## ---
## Signif. codes:  0 '***' 0.001 '**' 0.01 '*' 0.05 '.' 0.1 ' ' 1
##
## Residual standard error: 407200 on 10 degrees of freedom
## Multiple R-squared:  0.5161, Adjusted R-squared:  0.4677
## F-statistic: 10.67 on 1 and 10 DF,  p-value: 0.00849



### Kinetic of 500-m/z-associated compound
cor.test(mrm$Time_h, mrm$MRM_500, method = "spearman")

## Warning in cor.test.default(mrm$Time_h, mrm$MRM_500, method = "spearman"
## ):
## Cannot compute exact p-value with ties

##
##  Spearman's rank correlation rho
##
## data:  mrm$Time_h and mrm$MRM_500
## S = 49.239, p-value = 0.0008853
## alternative hypothesis: true rho is not equal to 0
## sample estimates:
##       rho
## 0.8278374

plot(mrm$MRM_500~log(mrm$Time_h))
abline(lm(mrm$MRM_500~log(mrm$Time_h)))
```

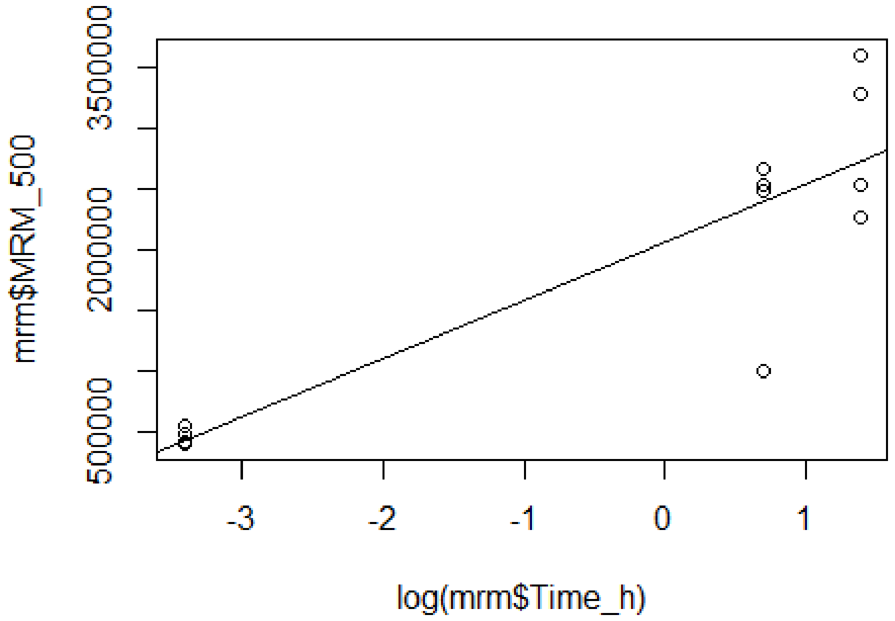


```
summary(lm(mrm$MRM_500~log(mrm$Time_h)))

##
## Call:
## lm(formula = mrm$MRM_500 ~ log(mrm$Time_h))
##
## Residuals:
##      Min       1Q   Median       3Q      Max
## -1403468   -65459    68471   166799   876463
##
## Coefficients:
##                 Estimate Std. Error t value Pr(>|t|)
## (Intercept)      2065723     172361  11.985 2.96e-07 ***
## log(mrm$Time_h)   480941      79677   6.036 0.000126 ***
## ---
## Signif. codes:  0 '***' 0.001 '**' 0.01 '*' 0.05 '.' 0.1 ' ' 1
##
## Residual standard error: 584400 on 10 degrees of freedom
## Multiple R-squared:  0.7846, Adjusted R-squared:  0.7631
## F-statistic: 36.43 on 1 and 10 DF,  p-value: 0.0001259
```
