## Supplemental File S3 for "Uncyclized xanthommatin is a key ommochrome intermediate in invertebrate coloration"

### Supplemental File S3 – Supplemental Discussion

#### The formation of decarboxylated xanthommatin

We propose three hypotheses to explain the presence of decarboxylated xanthommatin in biological samples. Here, we describe two of them, the third one being developed in the main text. (1) The decarboxylation of xanthommatin in water-based environments and upon light radiations possibly accounted for the formation of decarboxylated xanthommatin (Fig 8). Indeed, some aromatic compounds are known to be decarboxylated by the action of water (Mundle and Kluger, 2009), and kynurenic acid, which is structurally related to xanthommatin, is decarboxylated upon light radiations (Zelentsova et al., 2013). (2) Enzymatic and non-enzymatic syntheses of ommatins might differ in their products and exact molecular steps (Ishii et al., 1992; Zhuravlev et al., 2018). Thus, the proposed involvement of the heme peroxidase Cardinal in the biosynthesis of ommatins (Howells et al., 1977) might lead to the natural formation of xanthommatin over its decarboxylated form.

#### The biological function of decarboxylated xanthommatin

Few studies have addressed the biological function of decarboxylated xanthommatin. Its ratio to xanthommatin is known to vary among species and individuals (Futahashi et al., 2012; Williams et al., 2016). In cephalopods, the ratio of xanthommatin to its decarboxylated form within a chromatophore has been suggested to determine its color, ranging from yellow to purple (Williams et al., 2016). However, our absorbance data do not support this hypothesis because the experimental absorbance spectrum of decarboxylated xanthommatin was not different from that of xanthommatin in the visible region. Moreover, purple colors are produced by chromophores that absorb wavelengths around 520 nm, which has not been described for any ommatin in contrast to ommins (Figon and Casas, 2019). Another study focusing on the quantum chemistry of pirenoxine, a xanthommatin-like drug, proposed that the carboxylic acid function present on the pyrido[3,2-*a*]phenoxazinone of pirenoxine could enhance its binding to divalent cations (Liao et al., 2011). This is coherent with the fact that ommochromasomes accumulate and store cations, such as $Ca^{2+}$ and $Mg^{2+}$ (Gribakin et al., 1987; Ukhanov, 1991). Thus, favoring xanthommatin over its decarboxylated form *in vivo* might enhance the storage of metals in ommochromasomes, as proposed for eumelanins that contain high proportions of the carboxylated monomers DHICA (Hong and Simon, 2007). To which extent the binding of metals modifies the physical and chemical properties of ommatins, therefore their biological roles, remains to be determined.

#### The enzymatic formation of uncyclized xanthommatin

Whether the *in vivo* oxidative condensation of 3-hydroxykynurenine is catalyzed enzymatically remains a key question in the biogenesis of ommochromes (Figon and Casas, 2019). Theoretical calculations suggested that both enzymatic and non-enzymatic oxidations of 3-hydroxykynurenine would lead to the formation of the phenoxazinone uncyclized xanthommatin (Williams et al., 2019; Zhuravlev et al., 2018). There was some evidence of a phenoxazinone synthase (PHS) activity associated to purified ommochromasomes of fruitflies (Yamamoto et al., 1976). However, no corresponding PHS enzyme has ever been isolated nor identified in species producing ommochromes (Figon and Casas, 2019). PHS is not the only enzyme capable of forming aminophenoxazinones from *ortho*-aminophenols. Tyrosinase, laccase, peroxidase and catalase can also catalyze these reactions (Le Roes-Hill et al., 2009). Particularly, peroxidase can produce xanthommatin and its decarboxylated form from 3-hydroxykynurenine *in vitro* (Ishii et al., 1992; Iwahashi and Ishii, 1997; Vazquez et al., 2000; Vogliardi et al., 2004), likely through the formation of uncyclized xanthommatin (Iwahashi and Ishii, 1997). This result relates to the long-known fact that insects mutated for the heme peroxidase Cardinal accumulate 3-hydroxykynurenine without forming ommochromes (Howells et al., 1977; Osanai-Futahashi et al., 2016). Hence, our data support the hypothesis that the biosynthesis of ommatins could be catalyzed by a relatively unspecific peroxidase such as Cardinal (Fig 8) (Liu et al., 2017; Osanai-Futahashi et al., 2016), without the requirement of a specialized PHS.
